# Pyruvate Kinase M Links Glucose Availability to Protein Synthesis

**DOI:** 10.1101/715086

**Authors:** Nevraj S. Kejiou, Lena Ilan, Stefan Aigner, Enching Luo, Ines Rabano, Nishani Rajakulendran, Hamed S. Najafabadi, Stephane Angers, Gene W. Yeo, Alexander F. Palazzo

## Abstract

How human cells coordinate various metabolic processes, such as glycolysis and protein translation, remains unclear. One key insight is that various metabolic enzymes have been found to associate with mRNAs, however whether these enzymes regulate mRNA biology in response to changes in cellular metabolic state remains unknown. Here we report that the glycolytic enzyme, pyruvate kinase M (PKM), inhibits the translation of 7% of the transcriptome in response to elevated levels of glucose and pyruvate. Our data suggest that in the presence of glucose and pyruvate, PKM associates with ribosomes that are synthesizing stretches of polyacidic nascent polypeptides and stalls the elongation step of translation. PKM-regulated mRNAs encode proteins required for the cell cycle and may explain previous results linking PKM to cell cycle regulation. Our study uncovers an unappreciated link between glycolysis and the ribosome that likely coordinates the intake of glycolytic metabolites with the regulation of protein synthesis and the cell cycle.

## Results and Discussion

### Mass spectrometry analysis of ER and cytosolic polysomes and mRNPs

Over the past decade, numerous proteins that lack RNA binding domains, such as metabolic enzymes, have been shown to exhibit mRNA and ribosome binding^1–12^. It is unclear whether these unconventional RNA binding proteins (RBP) associate with transcripts and ribosomes in a spatially defined manner, or if they link metabolic states to mRNA stability or translation. To determine the spatial distribution of RNA-, and ribosome-binding proteins, we isolated ER and cytosolic fractions from human osteosarcoma (U2OS) cells (Figure 1A) and sedimented crude polysomes. The cytosol and ER represent the major division in cellular protein synthesis, each containing distinct pools of mRNAs and unique translational regulatory systems^13–16^. These isolated polysomes were then treated with RNase to liberate RNA-binding proteins (“RNA-bound fraction”), and resedimented to pellet ribosomes and associated proteins (“Ribosome-bound fraction”; Figure 1B). We then analyzed the composition of the RNA-bound and Ribosome-bound fractions (Figure 1B-C) by mass spectrometry, as previously described^17^. In parallel we also isolated messenger ribonuclear protein (mRNP) complexes from the ER and cytosol using oligo-dT affinity chromatography (“mRNP-bound”; Figure 1B, D, “dT”) and again analyzed these fractions by mass spectrometry. To control for non-specific binding to the oligo-dT resin we also performed the affinity chromatography with beads lacking any nucleic acid (Figure 1B, “Mock Beads”, 1D “B”). Our purification conditions, done in the absence of crosslinking, enabled recovery of proteins that are directly and indirectly bound to mRNAs and/or ribosomes.

**Figure 1.**
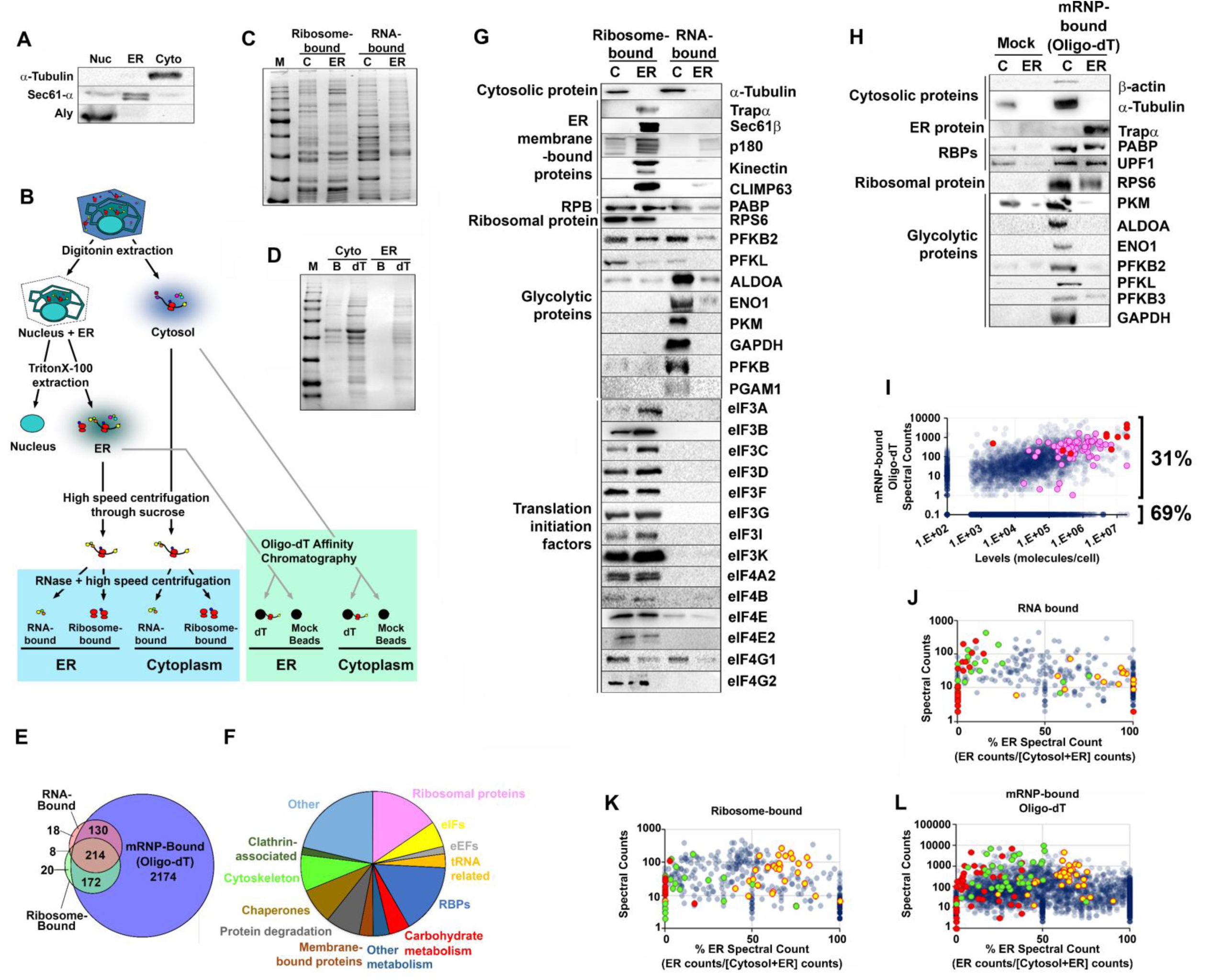
The distribution of different classes of proteins between cytosolic and ER polysomes. A) U2OS cell fractions probed for a cytosolic marker (α-tubulin), an ER-marker (Sec61α) and a nuclear marker (Aly). B) Schematic representation of the cell fractionation protocol. Crude polysome fractions are in the blue box, oligo-dT affinity chromatography fractions are in the green box. C-D) Coomassie stains of the various fractions (‘M’: molecular weight marker, ‘C’: cytosol, ‘B’: mock beads, ‘dT’: oligo-dT). E) The set of proteins enriched in either the two crude polysome fractions (RNA-associated: red, or ribosome-associated: green) or the oligo-dT associated fraction (blue) from U2OS cell fractions. F) The 496 proteins that were identified in both (1) the oligo-dT affinity purification and (2) one of the two crude-polysome fractions, divided up into different functional classes (see Supplementary Table 1 for the complete list). G-H) Analysis of the ER/cytosolic distribution of various proteins in the crude ribosome fractions (G) and oligo-dT affinity purified fractions (H) by immunoblot. I) For all proteins expressed in U2OS cells, a comparison of the total number of peptides identified by mass spectrometry in the oligo-dT chromatography experiments (both ER and cytosol; *y-axis*) was plotted against the estimated level of proteins (*x-axis*; data from ref 15). Glycolytic enzymes are labeled in red, and ribosomal proteins in pink. Proteins that did not appear in the oligo-dT chromatography experiments were set to 0.1 peptides, while those proteins that did not appear in the analysis of protein levels were set to 100 molecules/cell. J-L) For each protein present in either the crude polysome or oligo-dT purifications, the total number of peptides (*y-axis*) identified by mass spectrometry was plotted against the percentage of peptides found in the ER [100% x peptides in ER/(peptides in ER + peptides in cytoplasm); ‘% on ER’ - *x-axis*]. This analysis was performed on the RNA-bound (J) and ribosome-bound (K) fractions of the crude polysome preparations, and on the oligo-dT associated proteins (L). Note that the *x-axis* is the average of 5 biological replicates for (J-K) and 2 biological replicates for (L), while the *y-axis* is the total sum of peptides in all experiments (J-L). Classes of proteins that are enriched in either the ER or the cytoplasm are highlighted – carbohydrate metabolic proteins (red), eIFs (yellow), and cytoskeletal-associated proteins (green).

After statistical processing (see Methods), 370 proteins were present in the RNA-bound fraction, 414 were present in the ribosome-bound fraction, and 2690 proteins were enriched in the mRNP-associated fraction. Upon further manual curation (see methods), 496 proteins were identified in both (1) the oligo-dT-associated fraction and (2) one of the two crude polysome fractions (Figure 1E, Supplemental Table 1). The majority of these proteins (88%; 438/496) had been identified in previous global analyses of RNA- and ribosome-binding proteins^1–12^ (Supplemental Table 2), indicating that our list of polysome-interacting proteins is mostly in agreement with the literature.

The curated 496 mRNA/Ribosome-bound proteins fell into several broad categories (Figure 1F). This includes a wide array of metabolic enzymes (Figure 1F), especially those involved in carbohydrate metabolism, similar to what had been uncovered by several genome-wide RNA/Ribosome binding protein surveys from a variety of species^1–12, 18^. The most abundant by spectral counts were glycolytic enzymes, including both spliced isoforms of pyruvate kinase (PKM1 and 2). The detected glycolytic enzymes were not nascent polypeptides emerging from the ribosome, a concern for abundantly expressed proteins, as we found no bias for N-terminal peptides (Supplementary Figure 1), which would be the case for partially synthesized proteins. Furthermore, immunoblot analysis indicated that glycolytic enzymes were present in these fractions as full-length proteins (Figure 1G-H). Lastly, when the number of peptides spectral counts in the oligo-dT pulldown were compared to overall protein levels^19^, glycolytic enzymes were on par with ribosomal proteins (Figure 1I, glycolytic enzymes are in red, ribosomal proteins are in magenta), suggesting that they were not trace contaminants. The association of glycolytic enzymes with polysomes is largely lost upon RNase-treatment, (compare ribosome- and RNA-associated fractions, Figure 1G), indicating that these proteins bind to RNA either directly or through some intermediary protein.

### Certain classes of proteins are enriched in either cytosolic or ER-associated polysomes/mRNPs

The overall ER/cytosolic distribution of proteins for the RNA-, ribosome-, and mRNP-bound fractions were calculated by tabulating the normalized number of peptide spectral counts in the ER over the combined counts in the ER and cytosol (thus obtaining a percent ER peptide spectral count, Figure 1J-L, Supplemental Table 1). Using this analysis, we found that the ER/cytosolic distribution of proteins for each fraction differed significantly to what would be expected if peptides were randomly assorted (Supplemental Figure 2). In the ribosome-bound fractions, ribosomal proteins were found at comparable levels between the ER and cytosol (Supplemental Figure 3) suggesting that ribosomes were sampled equally from these two subcellular compartments. When the RNA-, ribosome-, and mRNP-bound fractions were evaluated by immunoblot, ER-resident membrane-bound proteins were consistently biased towards the ER, while poly(A)-binding protein and the ribosomal protein RPS6 were present at equivalent levels in the ER and cytosol (Figure 1G-H).

Interestingly, there were certain unexpected classes of proteins that had biased ER/cytosolic distributions in all three (RNA-, ribosome-, and mRNP-bound) fractions. Eukaryotic translation initiation factors (eIFs) were slightly enriched in the ER for all fractions (Figure 1J-L, yellow data points), and this was verified by immunoblot (Figure 1G). In contrast, cytoskeletal components including actin, tubulin, motor and cytoskeletal-binding proteins were enriched in the cytosol for all fractions (Figure 1J-L, green data points; also see α-tubulin in Figure 1G-H and β-actin in Figure 1H). Interestingly, carbohydrate metabolic enzymes, including glycolytic enzymes, were also enriched in the cytosol (Figure 1J-L, red data points). Indeed, most of these co-sedimented with cytosolic crude-polysomes in an RNA-dependent manner, suggesting that they bound directly or indirectly to RNA, and this was verified by immunoblot (Figure 1G-H).

### PKM associates with polysomes in the presence of glucose and pyruvate

Since we observed that glycolytic enzymes were present in cytosolic polysomes, we asked if their polysome-association was sensitive to changes in glycolysis, as the canonical functions of glycolytic enzymes are often regulated by metabolites. We found that PKM, which catalyzes the last step of glycolysis, preferentially co-fractionates with polysomes isolated from cells fed with a mixture of glucose and pyruvate in comparison to cells starved of these two metabolites for 3 hours (Figure 2A-C). When we isolated crude-polysomes by sedimentation through a sucrose cushion, PKM co-pelleted in a glucose/pyruvate-dependent manner (Figure 2D). In contrast, glucose/pyruvate-starvation did not change total cellular levels of PKM (Figure 2E) and did not significantly alter the levels of polysomes, ATP and lactic acid (Figure 2A-B, F-G), suggesting that this treatment did not activate major stress pathways in the cells. Moreover, this treatment only led to a slight decrease in mTOR activity, as assessed by the small drop in phosphorylated S6 kinase (Figure 2E), a downstream substrate of the mTOR kinase. In contrast, treatment with 2-deoxyglucose, a potent inhibitor of glycolysis, resulted in a drastic drop in phosphorylated S6 kinase, ATP and lactate (Figure 2E-G).

**Figure 2.**
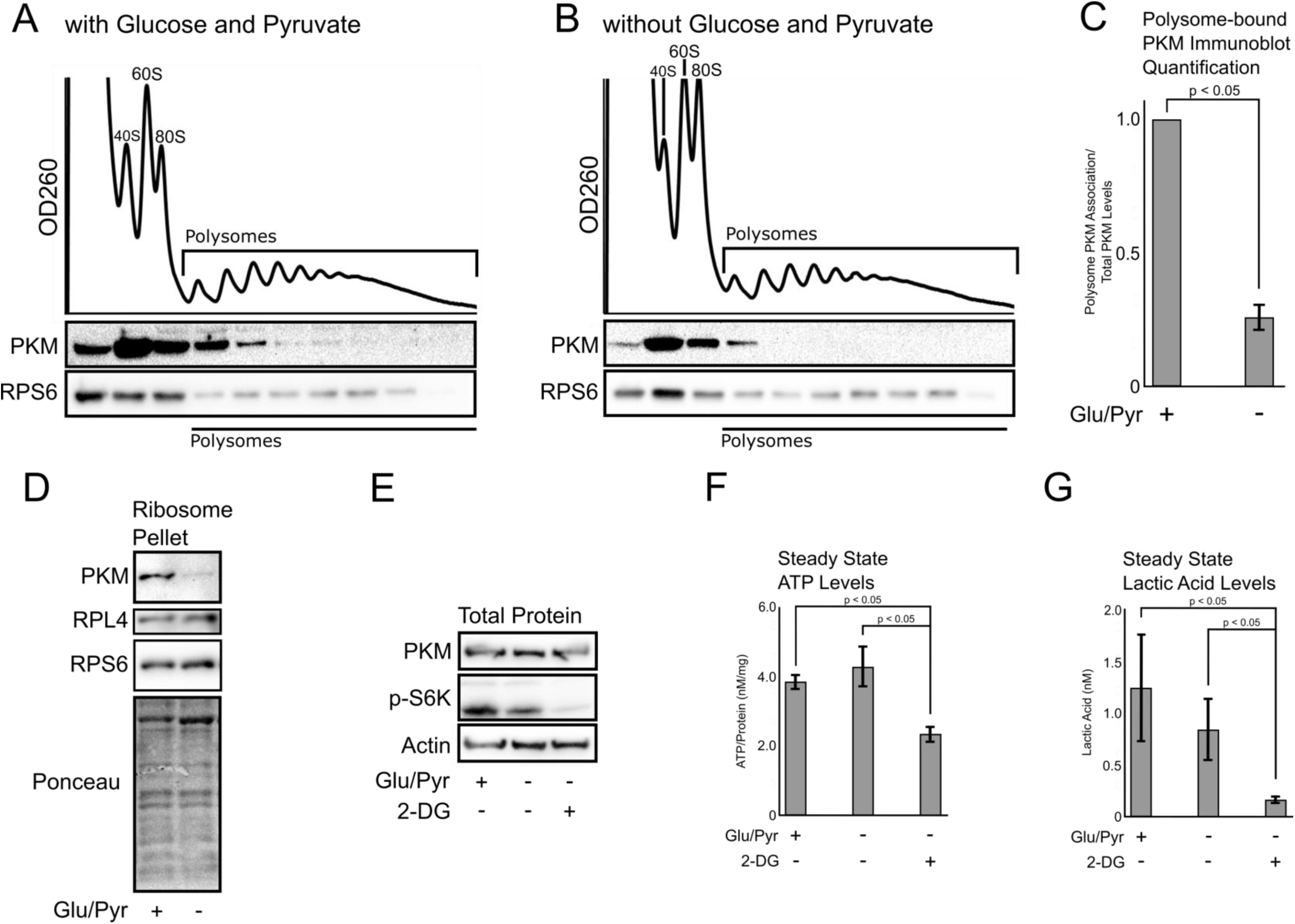
PKM associates with polysomes in a glucose/pyruvate-dependent manner. A-B) U2OS cells were incubated in DMEM that contained (A) or lacked (B) glucose (4.5 g/L) and/ sodium-pyruvate (0.11 g/L) for 3 hours. 10 minutes prior to lysis, the cells were treated with 100 μg/mL cycloheximide to stabilize polysomes. Cell lysates were sedimented through a 20-60% linear sucrose gradients at 160,000 *g* for 2 hours. Nucleic acids were monitored by OD260, and fractions were monitored by immunoblot for pan-PKM (both PKM1 and 2) and RPS6. C) Densitometry analysis of the fraction of PKM that associates with the polysome fractions (Figure 2A-B, lanes 4-10), normalized to total PKM (lanes 1-10). Each bar is the average and standard error of 3 biological replicates. D) U2OS cells were incubated in DMEM that contained or lacked glucose/pyruvate, with or without 2-deoxyglucose (‘2-DG’; 20 mM) for 3 hours. Lysates were monitored by immunoblot for pan-PKM, phosphorylated S6 Kinase (p-S6K, a downstream product of mTOR kinase) and Actin. E) U2OS cells were incubated in DMEM that contained or lacked glucose/pyruvate. Crude polysomes were sedimented as in Figure 1B and probed for pan-PKM and ribosomes (RPL4 and RPS6). Total proteins were also monitored by Ponceau stain. F-G) U2OS cells were incubated in DMEM that contained or lacked glucose/pyruvate, with or without 2-DG for 3 hr. Cell lysates were assessed for ATP (F) and Lactate (G) levels. Each bar is the average and standard error of 3 biological replicates.

### PKM associates with translating ribosomes

Given that we and others^1, 6, 10–12, 18^ have identified PKM as a putative RBP, we immunoprecipitated the major isoform, PKM2^20^ (Figure 3A), under very stringent conditions from lysates of UV-crosslinked HEPG2 cells, followed by limiting RNase treatment. When these immunoprecipitates were labeled with [γ^32^P]-ATP using polynucleotide kinase, one major RNase-sensitive band, corresponding to PKM2 was observed (Figure 3B), indicating that this protein was in close proximity to endogenous RNAs. Next, we isolated the cross-linked RNAs and analyzed these by eCLIP-Seq^21^. We identified approximately 4,000 enriched binding peaks in comparison to the size-matched inputs (8-fold enrichment, P-value <10^-5^), distributed over transcripts from 961 genes from two independent replicates (Supplemental Tables 3, 4). Strikingly, we found that PKM2 was crosslinked primarily to the coding sequence (CDS) of target transcripts (Figure 3C, D). This suggested that PKM2 is in close proximity to mRNA regions that are actively translated, and this may indicate that it interacts with the ribosome. This is in agreement with a previous study that identified PKM as a ribosome-associated protein^10^. Indeed, we observed that purified GST-PKM1 co-sedimented with salt-washed ribosomes in vitro (Supplemental Figure 5). In cell extracts, endogenous PKM co-sedimented with puromycin-treated ribosomes, which are not bound to mRNA and nascent chains; however, this association was disrupted by high salt (Figure 3E, F). PKM also co-sedimented with ribosomes from extracts treated with cycloheximide, which maintains ribosome-mRNA-nascent chain interactions, however under these conditions, PKM co-sedimentation was salt-insensitive (Figure 3E, F). This suggests that either PKM has a higher affinity for translating ribosomes, or that PKM forms contacts with the mRNA or nascent polypeptide chain. In contrast, poly-A binding protein (PABP) co-sedimented with ribosomes under all conditions (Figure 3E, F), consistent with previous findings^22, 23^. Since the binding of PKM to polysomes is partially RNase-sensitive (Figure 1G), it is possible that PKM recognizes rRNA, as suggested by previous work^10^, and our results indicate that PKM associates to crude polysomes in an RNA-dependent manner (Figure 1G). Indeed, we observe PKM2 eCLIP peaks on sections of mature rRNA (Supplemental Figure 6A-C); however, these do not align with previously-identified PKM2 CLIP reads, and overall rRNA was not enriched over the input (Supplemental Figure 6D).

**Figure 3.**
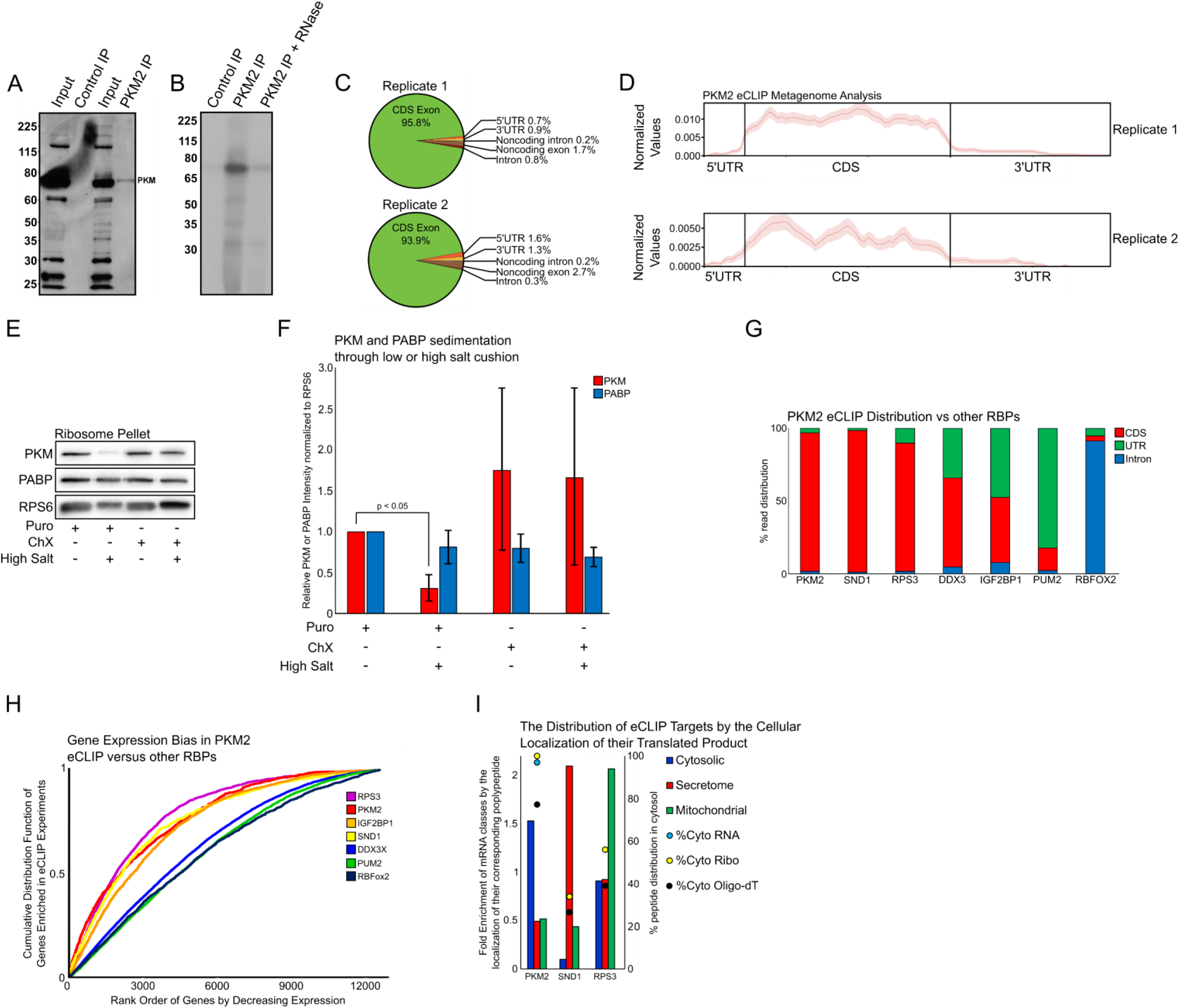
PKM2 associates with the coding sequence of mRNAs encoding cytosolic proteins. A) PKM2 was immunoprecipitated from HepG2 cell lysates, separated by SDS PAGE and immunoblotted. B) Mock (‘Control IP’) and PKM2-immunoprecipitates (‘PKM2 IP’) were labeled with γ^32^P]-ATP using polynucleotide kinase. To verify whether the band in question contained labeled RNA, the phosphorylated immunoprecipitate was treated with RNase (‘+ RNase’). The resulting reactions were resolved by SDS-PAGE and processed for autoradiography. One major band of ∼72kDa, corresponding to PKM2, was observed. C) Mapping of PKM2 eCLIP reads (8-fold enrichment, P-value <10^-5^) to various annotated parts of the human transcriptome. D) Mapping of PKM2 eCLIP reads along the normalized length of protein coding mRNAs. E-F) U2OS cells were treated with either 200 μM Puromycin (200 μM) or Cycloheximide (10 µg/mL) for 30 minutes and then lysed and incubated in isotonic (125 mM KCl) or a high salt (500 mM KCl) buffer. Ribosomes/polysomes were then isolated by centrifugation through a sucrose cushion and the pellets were analyzed by immunoblot for pan-PKM, PABP and RPS6 (E). PKM and PABP levels were monitored by densitometry analysis (F) with each bar representing the average and standard error of 3 biological replicates. G) The distribution of eCLIP reads for PKM2 and other RBPs (see Ref. 20) to annotated features of protein-coding genes. H) Cumulative distribution function demonstrating the bias of PKM for binding the mRNA of abundantly expressed genes. Rank order of genes by decreasing expression (*x-axis*) was plotted against their enrichment propensity in different RBP eCLIPs (*y-axis*, see Ref. 20). Note that RPS3, PKM2 IGF2BP1 and SND1 eCLIP targets are enriched for highly expressed mRNAs. In contrast, DDX3X, PUM2 and RBFox2 eCLIP targets do not have as much of a bias. I) For each RBP, their bound mRNAs identified by eCLIP were analyzed to determine what type of protein they encoded (enrichment *y-axis* on the left, “1” denotes no enrichment). In addition, the cytosolic distribution [100% x peptides in cytosol/(peptides in ER + peptides in cytosol); ‘% on ER’ - *y-axis* on the right] of each RBP in crude polysome and oligo-dT affinity fractions was plotted as in Figure 1J-L.

When the eCLIP-Seq with PKM2 was compared to similar experiments performed with other RBPs^24^, we observed that CDS-restricted RBPs, such as PKM2, SND1 and the RPS3 ribosomal protein (Figure 3G), were enriched for highly expressed mRNAs (Figure 3H). In contrast, most RBPs that display some binding to UTRs (DDX3X and PUM2) or introns (RBFOX2), did not show this bias (Figure 3G-H), with the exception of IGF2BP1. We assume that the CDS-restricted RBPs (PKM2, SND1 and RPS3) are ribosome-associated, while the others represent proteins that associate primarily with mature mRNAs and pre-mRNAs. Ribosome-associated RBPs may have this bias as highly expressed mRNAs tend to be more efficiently translated^25^. It is also worth noting that despite the fact that PKM2 and SND1 eCLIP targets were enriched for highly expressed mRNAs, the PKM2 eCLIP was enriched for mRNAs encoding cytosolic proteins, while SND1 eCLIP was enriched for mRNAs encoding proteins that are synthesized on the ER (“secretome”) (Figure 3I), and this reflects the enrichment of PKM in the cytosolic-derived mRNPs/polysomes and the enrichment of SND1 in the ER-derived mRNPs/polysomes (Figure 3I). In contrast, RPS3 eCLIP targets were not biased for the ER or cytosol, and RPS3 peptides were found in both fractions (Figure 3I).

### PKM promotes the polysome-association for a subset of mRNAs in response to glucose/pyruvate

Previously, it was found that PKM-depletion resulted in a decrease of ribosomal occupancy for a subset of mRNAs^10^. Given that PKM-binding to translating mRNA is glucose/pyruvate dependent (Figure 2A-C, E), we tested how these factors contributed to polysome-association over the whole transcriptome. We depleted PKM in U2OS cells by RNAi using lentiviral-delivered shRNAs (Figure 4A) and then either maintained the cells in normal or glucose/pyruvate-free medium for 3 hours. Overall, this brief glucose/pyruvate-starvation had minimal effects on overall polysome profiles (Figure 4B). We determined the degree of polysome-association for each mRNA by calculating the ratio of the abundance of mRNAs in the polysome (disomes or higher) fractions to the abundance in the total input by RNA-Seq (Supplemental Table 5). Strikingly, PKM-depletion decreased the polysome-association of a subset of mRNAs *only* in the presence of glucose/pyruvate (Figure 4C, Supplemental Figure 7A-B). This is to be expected as PKM predominantly associates with polysomes in the presence of glucose/pyruvate (Figure 2A-C, E). Moreover, we found that the short glucose/pyruvate starvation decreases the polysome-association of a subset of mRNAs in control cells, and that this effect was lost upon PKM-depletion (Figure 4D, Supplemental Figure 7C), indicating that the regulation of polysome-association was dependent on the presence of PKM. PKM-depletion had minimal effects on steady state levels of most mRNAs (Supplemental Figure 8). When we performed a three-way comparison between mRNAs enriched in the PKM2 eCLIP, to mRNAs whose polysome-association was dependent on either glucose/pyruvate or PKM, we observed that all three sets significantly overlapped (Fig. 4E, F). Again, mRNAs whose polysome-association was glucose/pyruvate- and PKM-regulated were enriched for those encoding cytosolic proteins (Figure 4G).

**Figure 4.**
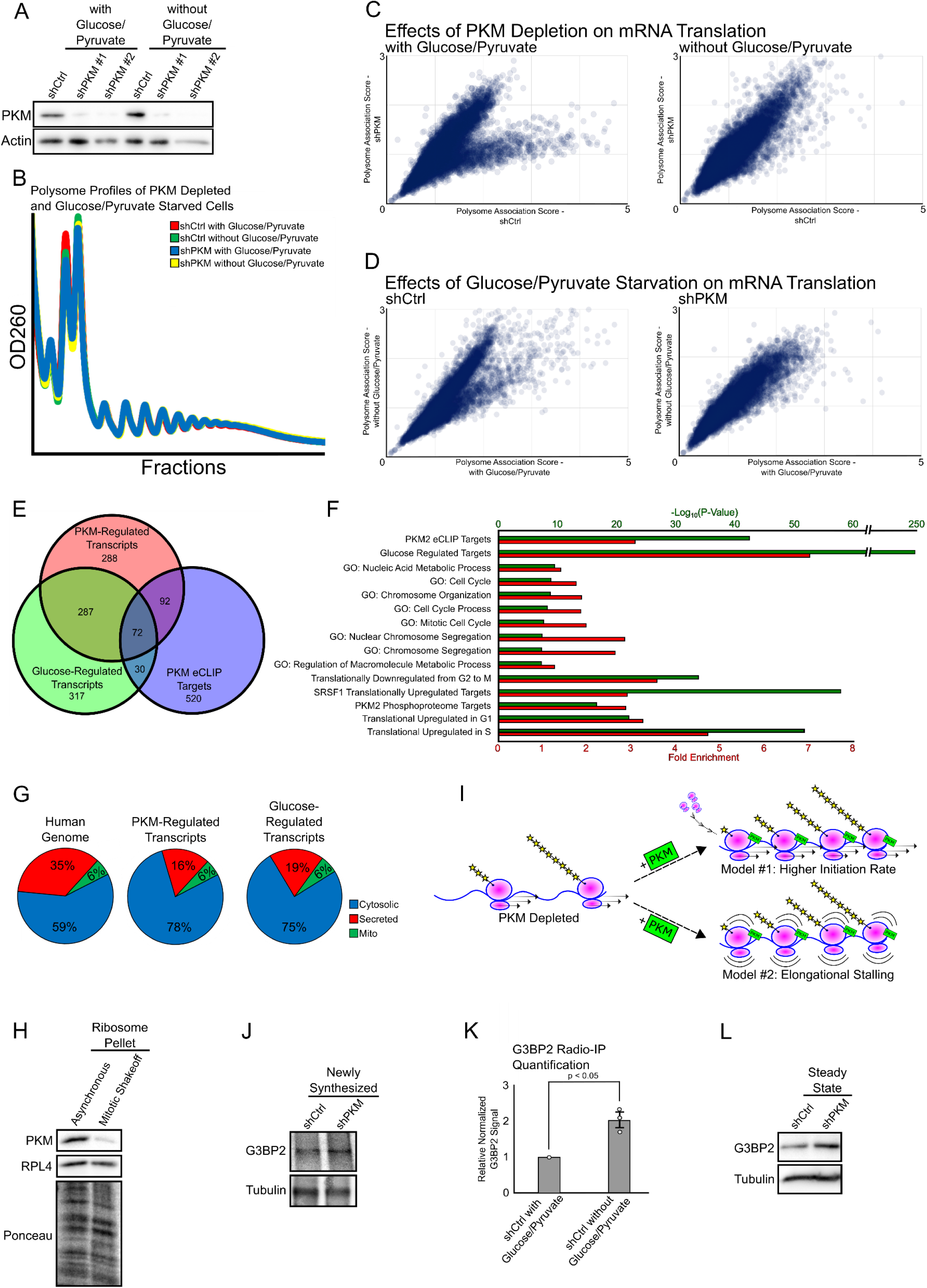
PKM inhibits the translation elongation of ribosomes on a subset of mRNAs in a glucose/pyruvate-dependent manner. A-B) PKM was depleted from U2OS cells using lentiviral-delivered shRNAs. Four days post-infection, cells were either incubated in DMEM with glucose/pyruvate or glucose/pyruvate-free DMEM for 3 hours. 30 minutes prior to lysis, the cells were treated with 100 μg/mL Lysates were either separated by SDS PAGE and immunoblotted for PKM and actin (A) or loaded onto a sucrose gradient, centrifuged, fractionated and analyzed by OD260 (B). C-D) RNA-Seq was performed on RNA isolated from total lysate or polysome fractions of the four samples shown in Figure 4B. The enrichment of each mRNA species in the polysome-associated fractions was plotted comparing transcripts isolated from PKM-depleted with control depleted cells (C) or comparing transcripts isolated from cells grown in normal and glucose/pyruvate-free media (D). Each dot is the average of 3 biological replicates. E) Three-way comparison of transcripts that were isolated in the PKM2 eCLIP experiment, to those that had a statistically significant increase (see Supplemental Table 5 and methods for further details) in their polysome-association in response to glucose/pyruvate-starvation treatment or PKM-depletion. F) The set of mRNAs whose translation was sensitive to PKM-depletion was analyzed for significant overlap (red bars: fold enrichment – “1” denotes no enrichment, green bars: hypergeometric p-value) with other mRNA sets, gene ontology terms, and sets of mRNAs whose translation is downregulated from G2 to M^31^, whose translation is upregulated by SRSF1 overexpression^33^, whose translational products have been documented as substrates of PKM-dependent phosphorylation^39^, and whose translation is either upregulated in G1 or S^32^. G) For each mRNA set, the types of encoded proteins were analyzed to determined. Like PKM2 eCLIP targets (see Figure 3I), mRNAs whose translation was sensitive to glucose/pyruvate and PKM-depletion are enriched for those encoding cytosolic proteins. H) Normal asynchronous, or mitotically-arrested (double thymidine + nocodazole-treatment) U2OS cells were lysed, and crude polysomes were sedimented as in Figure 1B. Sedimented fractions were probed for pan-PKM and RPL4. Total proteins were also monitored by Ponceau stain. I) Two models for how glucose/pyruvate and PKM induce an increase in polysome-association. PKM either promotes translation initiation, or inhibits translation elongation. J-K) Newly synthesized proteins were labeled by feeding control or PKM-depleted U2OS cells with ^35^S-methionine/cysteine for 15 minutes, and G3BP2 and α-tubulin were immunoprecipitated. The newly synthesized proteins were separated by SDS-PAGE, imaged by auto-radiography (I), and levels were analyzed by densitometry analysis (J). Each bar is the average and standard error of 3 biological replicates. L) Total levels of G3BP2 and α-tubulin in control and PKM-depleted cell lysates as determined by immunoblot.

Interestingly, PKM-regulated mRNAs were enriched for GO terms that are related to cell cycle and were highly correlated with mRNAs that have been found to be differentially translated at different cell cycle stages (Figure 4F). Indeed, we found that PKM binding to polysomes decreased in mitotically-arrested cells (Figure 4H). This data suggests that PKM-association may affect ribosome activity only at certain phases of the cell cycle.

### PKM stalls translational elongation

An increase in polysome-association or ribosome footprints on a given mRNA may suggest either an increase in translation initiation or stalling during translational elongation^26, 27^ (Figure 4I). The former possibility would lead to an increase in protein production, while the latter would result in a decrease. To distinguish between these two possibilities, we assessed PKM-depleted cells for newly made G3BP2 protein, whose mRNA was enriched in the PKM2 eCLIP, and also associated with polysomes in a PKM-and glucose-dependent manner. We thus labelled newly synthesized proteins using a pulse of ^35^S-methionine/cysteine for 15 minutes and immunoprecipitated G3BP2. We found that PKM-depletion led to an increase in newly synthesized G3BP2 (Figure 4J-K), consistent with the model that PKM interferes with translational elongation. In agreement with this finding, PKM-depletion upregulated overall G3BP2 levels (Figure 4L).

### PKM associates with ribosomes that synthesize charged nascent peptides

Given that PKM likely stalls elongating ribosomes, we asked how it recognizes specific targets. We found a motif that was enriched in the coding sequence of not only PKM2 eCLIP targets but also of mRNAs whose translation was inhibited by PKM and glucose/pyruvate (Figure 5A). Strikingly, this motif was *only* enriched in one particular reading frame, which encodes a short stretch of negatively charged amino acids, mostly aspartate. This motif was present mostly upstream (ranging from −180 to +40 nucleotides) of the center of PKM2 eCLIP peaks. If we assume that PKM2 crosslinks to mRNA near regions that are actively being decoded, then our results are consistent with PKM recognizing ribosomes that contain nascent polypeptide chains that are rich in negative amino acids. To further explore the link between PKM binding and the charge of the nascent polypeptide, we aligned the eCLIP peaks, converted these sequences to their cognate nascent polypeptide chains, and analyzed the distribution of charged amino acids (Figure 5C). Consistent with our motif analysis, we found that negative amino acids are enriched just upstream of the PKM2-eCLIP peaks. Again, if we assume that PKM crosslinks to mRNA regions that are undergoing translation, then the stretches of negative polypeptide would concentrate within the ribosome exit tunnel which accommodates about 30 amino acids (Figure 5D, highlighted window). To our surprise, we also observed that positive amino acids were concentrated in this window as well (Figure 5D). We then further characterized the encoded polypeptide from mRNAs that had both PKIM2-eCLIP peaks and whose translation was inhibited by either glucose/pyruvate or PKM. Interestingly, mRNAs whose translation is inhibited by glucose/pyruvate tended to code for negative but not positive polypeptide sequences near the eCLIP peak (Figure 5E). We saw less of a significant signal for PKM-regulated mRNAs (Figure 5F), however this may be due to the low number of transcripts in our analysis. Looking at individual targets, such as G3BP2, we could readily detect that the main eCLIP peak was downstream of a region that encodes an acidic polypeptide (Figure 5G).

**Figure 5.**
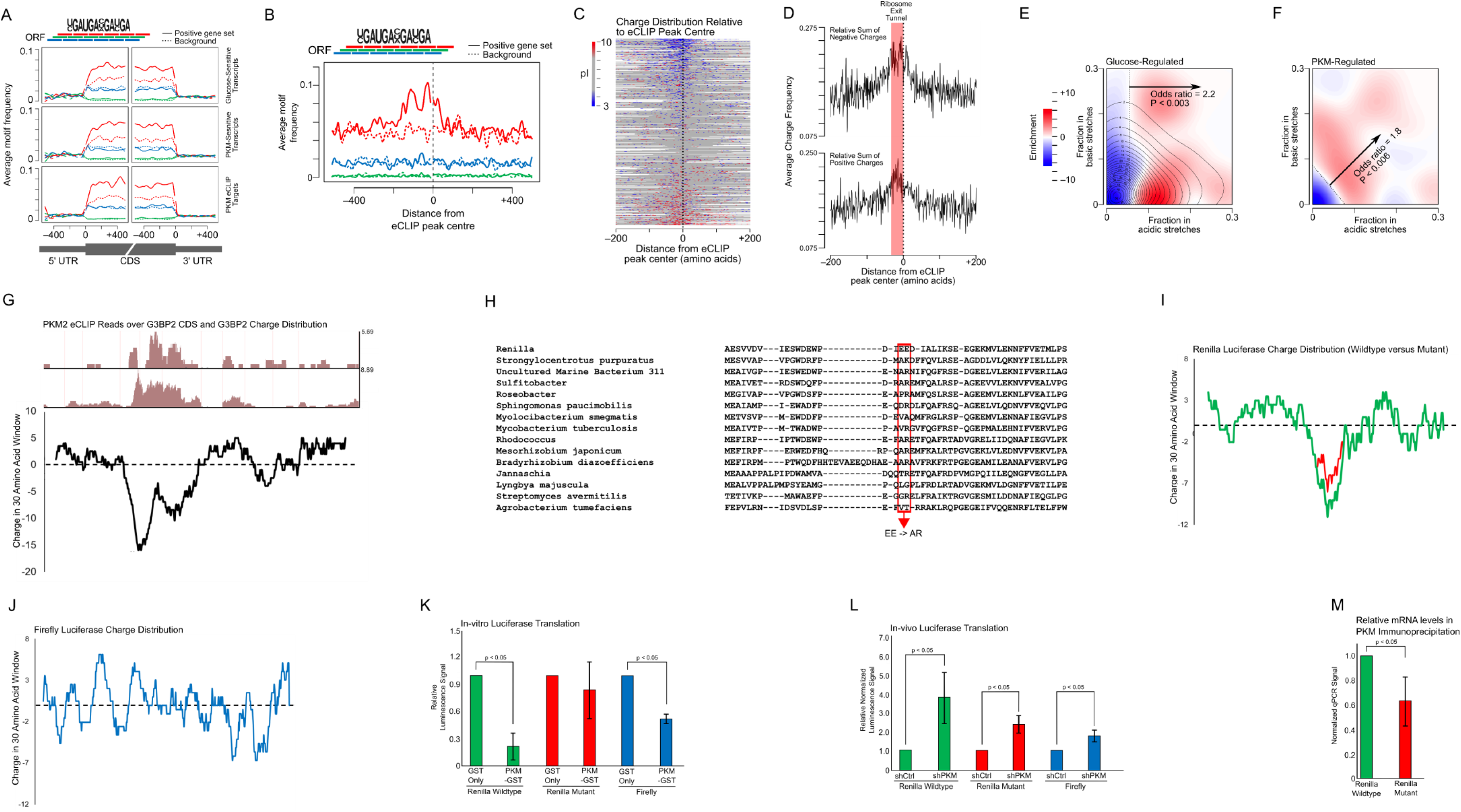
PKM inhibits the translation of mRNAs that encode polyacidic polypeptide stretches. A) Presence of an enriched motif, discovered by HOMER analysis, in various mRNA sets. Note that the motif is enriched in the open reading frame, and is only present in the red frame but not the blue or green ones. B) Distribution of the enriched motif in PKM2 eCLIP targets, with respect to the center of PKM2 eCLIP peaks (aligned at position “0” with dashed line). C) PKM2 eCLIP peaks were aligned (denoted at position “0” with dashed line) and the encoded polypeptides were analyzed for the presence of positive (blue) or negative (red) amino acids. Neutral amino acids are in grey. D) The frequency of negative and positive amino acids with respect to the center of PKM2 eCLIP peaks (denoted at position “0”, with dashed line). The amino acids corresponding to the 30 codons upstream of the PKM2 eCLIP peaks are highlighted in red. If we assume that each peak corresponds to PKM crosslinking to the mRNA region that is actively decoded, then the highlighted upstream 30 amino acids would be present in the exit tunnel of the ribosome. E-F) mRNAs that are translationally regulated by glucose/pyruvate (E) or PKM (F), and had PKM2 eCLIP peaks were analyzed for their decoded amino acid content in the regions surrounding the eCLIP peak (with a window of 200 amino acids upstream and downstream from each peak). Enrichment of either positive or negative charged amino acids were plotted. G) Comparison of PKM2 eCLIP peaks that map to the G3BP2 coding sequence, from UCSC genome browser track (https://s3-us-west-1.amazonaws.com/sauron-yeo/20170112_PKM2_CLIP/hub.txt), to the moving average of charge (in a 30 amino acid window) along the length of G3BP2 polypeptide sequence. H) Alignment of the luciferase polypeptide sequences from various organisms – centered around a poorly conserved stretch of negatively charged amino acids. Highlighted is a poorly conserved pair of glutamates that was mutated to alanine and arginine, which is found in many homologues. I-J) Plots of the moving average of charge (in a 30 amino acid window) along the length of the wildtype (green) and mutated (red) *Renilla* luciferase protein (I) and of the wildtype Firefly luciferase protein (J). K) In vitro synthesized Renilla and firefly *luciferase* mRNAs were translated using rabbit reticulocyte lysate supplemented with either GST or PKM1-GST. Levels of luminescence were assessed and normalized to the GST supplemented lysate. Each bar is the average and standard error of 3 biological replicates. L) Plasmids containing both Renilla and firefly *luciferase* genes were transfected into control or PKM-depleted U2OS cells. Levels of luminescence were assessed and normalized to the control cells 24 hours-post transfection. Each bar is the average and standard error of 5 biological replicates. M) Plasmids containing both Renilla and firefly *luciferase* genes were transfected into U2OS cells. 24 hours later, PKM was immunoprecipitated and the levels of Renilla (wildtype and mutant) and firefly *luciferase* mRNA were assessed by RT-qPCR. Levels of mRNA enrichment in the PKM immunoprecipitated over control precipitates were assessed and then normalized to the levels of firefly *luciferase* mRNA. Each bar is the average and standard error of 3 biological replicates.

### PKM inhibits the translation of mRNAs encoding polyacidic nascent peptide tracts

To test the effects of PKM on the production of proteins that contain acidic stretches, we monitored the translation of reporter luciferases in vivo and in vitro. Renilla luciferase protein contains one major polyacidic tract in a poorly conserved region (Figure 5H-I), while Firefly luciferase contains multiple small polyacidic amino acid tracts (Figure 5J). To test whether the encoded polyacidic tract affected the translation of the Renilla *luciferase* mRNA, we mutated two codons in the middle of the poorly conserved region so that they now encode non-acidic residues that are present in other luciferase homologues (Figure 5H-I). Using a rabbit reticulocyte in vitro translation system, we found that the addition of exogenous PKM1-GST decreased the translation of both Renilla and Firefly *luciferase* mRNAs, but not the mutant Renilla *luciferase* mRNA (Figure 5K). When plasmids of these various luciferase genes were transfected into U2OS cells, depletion of PKM led to an increase in the production of Renilla luciferase, and a smaller increase in the production of both the mutant Renilla and the Firefly luciferases (Figure 5L). Lastly, by performing a PKM2 RNA-immunoprecipitation, we found that PKM2 bound to Renilla *luciferase* mRNA (7.8+/- 3.1 fold enrichment in the anti-PKM versus control immunoprecipitation) and this association decreased for the mutant Renilla *luciferase* mRNA (Figure 5M).

### PKM, a regulator of mRNA translation

Overall, our data suggests that mRNAs encoding polyacidic polypeptide tracts are targets for PKM-mediated translational stalling in the presence of glucose/pyruvate. To our knowledge we provide the first transcriptome-wide analysis of glucose-regulated translation in mammalian cells.

In budding yeast, brief glucose-starvation drastically inhibits global translation^28–30^. In humans, this is not the case. Instead, glucose/pyruvate and PKM inhibit the translation of a small subset of mRNAs which encode proteins that are associated with cell-cycle and mitotic progression (Figure 4F). These same mRNAs were previously shown to have lower ribosome occupancy during M phase^31^ and higher occupancy during G1/S phase^32^ (Figure 4F), which could be due to PKM. This is consistent with the decrease in polysome-association of PKM in mitotically-arrested cells (Figure 4H). In addition to this, we also found a high correlation between PKM-regulated mRNAs and transcripts whose translation is enhanced by the over-expression of the RNA-binding protein SRSF1^33^(Figure 4F), which is upregulated in various cancers^34^. PKM-depletion has previously been reported to affect cell-cycle progression, predominantly through the loss of PKM-dependent non-canonical protein kinase activity or changes in metabolic activity^35–38^. Our new data suggests that PKM-mediated translational repression may also contribute.

Interestingly, we also find a significant correlation between mRNAs whose translation is regulated by PKM and mRNAs that encode for proteins that are targets of non-canonical PKM-phosphorylation^39^ (Figure 4F). Moreover, these phosphorylated serine and threonine residues tend to be surrounded by polyacidic polypeptide tracts. This observation may indicate that PKM binds to polyacidic nascent polypeptides, as suggested by our findings. It remains unclear whether PKM also affects the translation of mRNAs that contain polybasic polypeptide tracts as suggested by the eCLIP data (Figure 5D).

The role of PKM in cancer development is widely studied and our new findings provide a fresh lens to interpret old literature and a new foundation to understand how PKM regulates eukaryotic mRNA translation.

## Materials and Methods

### Cell types and growth conditions

U2OS cells were maintained in DMEM supplemented with 10% Fetal Bovine Serum (FBS) and 1% penicillin/streptomycin at 37°C and 5% CO_2_. Short term glucose starvation was carried out via incubation with glucose- and pyruvate-free DMEM (Gibco Cat#11966-025) supplemented with 10% FBS and 1% P/S with either 20 mM 2-deoxyglucose (D8375 Sigma) or vehicle (dH_2_O) for 3 hours.

### Cell fractionation and oligo-dT affinity chromatography

To isolate crude polysomal fractions, 75×10^6^ U2OS cells were treated with growth medium supplemented with 10 µg/mL cycloheximide for 30 minutes. Cells were collected by trypsinization, then pelleted at 800 *g* for 2 min, washed twice with ice cold PBS containing 10 µg/mL cycloheximide, washed once with ice cold Phy Buffer (150 mM Potassium Acetate, 5 mM Magnesium Acetate, 20 mM HEPES-KOH pH 7.4, 5 mM DTT, protease inhibitor cocktail [Roche], 10 µg/mL cycloheximide). Cell pellets were then re-suspended in 1 mL Phy Buffer. Cells were extracted by adding an equal volume of cold Phy Buffer + 0.08% digitonin and gently inverting the tube to allow the detergent to mix. The solution was then centrifuged at 800 *g* for 2 minutes to produce a suspension (cytosolic fraction, C1) and pellet (P1). The pellet was resuspended in additional 1 mL of cold Phy Buffer and extracted with an equal volume of cold Phy Buffer + 0.08% digitonin. The solution was then centrifuged at 800 *g* for 2 minutes to produce a suspension (C2) and pellet (ER + nuclear fraction, P2). The pellet was then resuspended in 1 mL cold Phy Buffer and extracted by adding an equal volume (1 mL) of Phy Buffer + 0.05% TritonX-100. This sample was then centrifuged at 800 *g* for 2 minutes to produce a suspension (ER fraction) and pellet (nuclear fraction). Cytosolic (C1) and ER fractions were then centrifuged at 10,000 *g* for 10 minutes to remove contaminating organelles such as mitochondria and nuclei. These fractions were analyzed for protein or further fractionated.

For crude polysome fractionation, 0.5ml of the C1 or P2 fractions were layered over a 0.5 ml Phy-Sucrose buffer (Phy buffer containing 1 M Sucrose), and centrifuged at 90,000 RPM for 40 minutes in a TLA120.2 rotor, to produce a suspension (non-polysomes) and a pellet (polysomes). The polysome fraction was then resuspended in 100 µL of Phy Buffer supplemented with 10 µL RNases and incubated for 10 minutes at room temperature. Treated fractions were centrifuged 60,000 rpm for 1 hour to produce a suspension (mRNA-bound proteins) and pellet (Ribosome-bound proteins) (see Figure 1A).

For the oligo-dT affinity chromatography 0.5 ml of C1 or P2 fractions were incubated with an equal volume of a 50% slurry of oligo(dT) beads (NEB #S1408S) in Phy buffer, or unconjugated Protein A beads (Thermo Fisher Cat#101041) overnight at 4□c Beads were washed 5 times with 1 mL cold Phy buffer. Proteins were eluted off the beads by incubating them with 2X Laemmli buffer at 65°C for 5 min.

### Polysome Profiling

18×10^6^ U2OS were treated with cycloheximide (100 μg/mL) for 10 minutes. Cells were collected via μ trypsinization and pelleted at 800 *g* and washed with ice cold PBS (supplemented with 100 μg/mL cycloheximide) three times. Cells were resuspended in 1 mL polysome lysis buffer (20 mM HEPES-KOH pH 7.4, 5 mM MgCl_2_, 100 mM KCl, 1% Triton X-100, 100 μg/ml cycloheximide) supplemented with 20U/mL Superase RNAse Inhibitor (Thermo Fisher Cat# AM2694), and protease inhibitor cocktail (Roche). Lysates were cleared via centrifugation at 16,000 *g* for 10 minutes. 500 μL of lysate saved for RNA input or total. The remaining 500 μL was layered on a 20% to 60% sucrose gradient buffer (20 mM HEPES-KOH pH 7.4, 5 mM MgCl_2_, 100 mM KCl, 100 μg/ml cycloheximide, and either 20% or 60% sucrose weight/volume) generated using the Biocomp Gradient Master. Lysates centrifuged at 36,000 *g* for 2 hours in a SW-41 rotor. Samples were collected and OD260 was continuously measured using the Biocomp Piston Gradient Fractionator. For RNA-seq analysis, polysome fractions (disomes and heavier) were pooled together and RNA was extracted using Trizol-LS (Thermofisher Cat#10296010) protocol. For total fraction, RNA was extracted from saved input. Forimmunoblotting, proteins from individual fractions were salted out via a TCA precipitation, washed in 100% acetone and re-suspended in 5X laemmli buffer.

### Ribosome co-sedimentation

18×10^6^ U2OS cells were pretreated with either cycloheximide (10 µg/mL) or puromycin (200 µM) for 30 minutes and then collected by trypsinization. The cells were washed twice in ice cold PBS and lysed in ribosome lysis buffer (125 mM KCl, 5 mM MgCl_2_, 20 mM HEPES-KOH pH 7.4, 250 mM sucrose, 0.08% Digitonin, 100 µg/mL cycloheximide). Unlysed cells were removed by centrifugation at 800 *g* for 10 minutes, and the resultant supernatant was centrifuged for 16,000 *g* to remove cellular debris. The concentration of KCl in the cleared supernatant was adjusted to 500 mM for high salt conditions or remained at 125 mM for low salt conditions. 0.5 ml of cleared supernatant was layered on 0.5 ml of either a high salt (500 mM KCl) or low salt (125 mM KCl) sucrose cushion (1 M sucrose, 5 mM MgCl_2,_ 20 mM HEPES-KOH pH 7.4, 100 µg/mL cycloheximide) in a 1 mL polycarbonate tube and then centrifuged at 90,000 RPM for 1 hour in a TLA-120.2 rotor. The pellet was washed twice in ice cold dH_2_O prior to solubilization in suspension buffer (125 mM KCl, 5 mM MgCl_2,_ 20 mM HEPES-KOH pH 7.4).

### Cell Synchronization and Mitotic Shakeoff

Approximately 8.8×10^6^ U2OS cells were synchronized toward G1/S by growing in medium supplemented with 2 mM thymidine for 16 hours, followed by 24 µM deoxycytidine for 8 hours, followed by 2 mM thymidine for 16 hours, and lastly by 24 µM deoxycytidine for 2 hours. 100 ng/mL of nocodazole was added for 18 hours to mitotically arrest synchronized cells. Mitotic cells were collected by vigorously washing tissue culture dish with PBS.

### Salt-Washed Ribosome Isolation

To generate salt washed ribosomes, 500 mL of HEK293F cells were collected via centrifugation at 800 *g*, and washed 5 times with ice cold PBS. Cells were lysed in modified ribosome lysis buffer (125 mM KCl, 5 mM MgCl_2_, 50 mM Tris-HCl pH 7.4, 250 mM sucrose, 1% NP-40). Lysates were cleared at 16,000 *g* for 10 minutes. The concentration of KCl in the cleared supernatant was adjusted to 500 mM KCl. 0.5 ml of cleared supernatant was layered on 0.5 ml of high salt (500 mM KCl) sucrose cushion (1 M sucrose, 5 mM MgCl_2,_ 50 mM Tris-HCl pH 7.4) in a 1 mL polycarbonate tube and then centrifuged at 90,000 rpm for 1 hour in a TLA-120.2 rotor. Pellet were re-suspended in modified suspension buffer (500 mM KCl, 5 mM MgCl_2,_ 50 mM Tris-HCl pH 7.4). Re-suspended pellets were subjected to an additional round of centrifugation – layering 0.5 mL of the re-suspended pellet solution over 0.5 mL of high salt cushion buffer in a 1 mL polycarbonate tube and then centrifuged at 90,000 RPM for 1 hour in a TLA-120.2 rotor. Pellets were then re-suspended in suspension buffer (25 mM KCl, 5 mM MgCl_2,_ 50 mM Tris-HCl pH 7.4). BCA assay was used to measure relative ribosome concentrations.

### In vitro Co-sedimentation

Equal amounts of salt-washed ribosomes, as isolated above, were mixed with equal molar amounts of either recombinant GST or GST-PKM1 in a 0.1 mL solution and incubated on ice for 30 minutes. This binding solution was layered on either a low salt (100 mM KCl) or high salt (200 mM KCl) 0.5 mL sucrose cushion (1 M sucrose, 5 mM MgCl_2,_ 50 mM Tris-HCl pH 7.4) in a 1 mL polycarbonate tube and then centrifuged at 90,000 RPM for 1 hour in a TLA-120.2 rotor. The supernatant was discarded and the pellet washed twice in ice-cold water. Pellet was re-suspended in 1X Laemmli buffer and denatured at 95°C for 5 min.

### In vitro Transcription

Renilla *luciferase* mRNA variants, both wildtypes and mutant, were generated from linearized pcDNA3 containing their respective coding sequences. In vitro transcription was carried as per HiScribe™ T7 Quick High Yield RNA Synthesis Kit (Cat#E2050S). Transcripts were cleaned as per PureLink RNA Mini Kit protocol (Cat#12183020). Transcripts were polyadenylated as per E. coli Poly(A) Polymerase Kit (Cat#M0276S) and was cleaned again using the PureLink RNA Mini Kit.

### In vitro Translation and Luciferase Assay

To perform in vitro *luciferase* translation assay, reactions were prepared as per Rabbit Reticulocyte Lysate System (Promega Cat#L4960). 10 µL of reaction mixture was supplemented with 1 µg of either Renilla or Firefly luciferase mRNA alongside recombinant GST or GST-PKM1. Renilla mRNA variants (either wildtype or mutant) was obtained via in vitro transcription, whereas Control Firefly mRNA was provided with Rabbit Reticulocyte Lysate System (Promega Cat#L4960). In vitro reaction was translated for 15 minutes at 37°C and stopped with the addition of 100 µg of cycloheximide and stored on ice prior to luciferase assay.

### Luciferase Assay

Both in vitro and in vivo luciferase assays were performed as per Dual Luciferase Reporter Assay System (Promega Cat#E1910) to obtain luminescence values for both Renilla and Firefly. Luminescence values were measured using the Berthold Sirius Single Tube Luminometer.

### Radio Immunoprecipitation

8.8×10^6^ U2OS cells were pulsed with 25 µCi of Cys/Met-protein label (Perkin-Elmer Cat#NEG772002MC) for 15 minutes. Cells were lysed using 1 m L radio-IP lysis buffer (20 mM HEPES-KOH pH7.4, 150 mM NaCl, 100 mM Iodoacetamide, protease inhibitor tablet, 1% NP-40). Lysate was cleared at 16,000 *g* for 10 minutes. Cleared lysates were incubated with either 4 µg of anti-Tubulin^40^ or anti-G3BP2 (Proteintech Cat#16276-1-AP) antibody for 30 minutes. 50 µL of protein-A slurry was added to lysates and were shaken overnight at 4°C. After immunoprecipitation, protein-A beads were washed 4 times using radio-IP wash buffer (20 mM HEPES-KOH pH7.4, 150 mM NaCl, 1% NP-40). IP samples were denatured, on beads, by the addition of 50 µL 5X Laemmli buffer and by heating at 95°C for 5 minutes. Samples were run through a 10% SDS-PAGE. Gel was fixed via a 15-minute incubation in fixing solution (50% MeOH, 10% Acetic Acid, 40% dH_2_O). Fixed gel was dried on Wattman paper. Radioactive gel was exposed, on film, for 1 week. Images were taken with a Typhoon-FLA 9000 imager.

### Immunoblot analysis

Samples were denatured with Laemmli sample buffer at 65°C for 5 minutes and separated by SDS-PAGE on 6 to 15% acrylamide gels. Separated proteins were transferred to nitrocellulose, stained by ponceau for general quality control and immunoprobed with antibodies as following: Aldolase-A (rabbit polyclonal, 1:1000, Cell Signaling Technology), Aly (rabbit polyclonal^41^, 1:1000, Sigma), α-Tubulin (mouse monoclonal DM1A, 1:5000, Sigma), β-Actin (mouse monoclonal, 1:1000, Sigma), eIF3A (rabbit β polyclonal, 1:1000, Novus), eIF3B (rabbit polyclonal, 1:1000, Bethyl), eIF3C (rabbit polyclonal, 1:1000, Bethyl), eIF3D (rabbit polyclonal, 1;1000, Bethyl), eIF3E (rabbit polyclonal, 1:1000, Bethyl), eIF3F (rabbit polyclonal, 1:1000, Bethyl), eIF3G (rabbit polyclonal, 1:1000, Bethyl), eIF3I (mouse monoclonal, 1:1000, Biolegend), eIF3K (rabbit polyclonal, 1:1000, Bethyl), eIF4A2 (rabbit polyclonal, 1:1000, Abcam), eIF4B (rabbit polyclonal, 1:1000, Signalway Antibody), eIF4E (rabbit polyclonal, 1:1000, Signalway Antibody), eIF4E2 (mouse monoclonal, 1:1000, Santa Cruz Antibody), eIF4G1 (rabbit polyclonal, 1:1000, Cell Signaling Technology), eIFG2 (rabbit polyclonal, 1:1000, Cell Signaling Technology), Enolase-1 (rabbit polyclonal, 1:1000, Cell Signaling Technology), G3BP1 (rabbit polyclonal, 1:1000, Signalway Antibody), G3BP2 (rabbit polyclonal, 1:1000, Proteintech), GAPDH (rabbit polyclonal, 1:1000, Abgent), Kinectin (rabbit polyclonal, 1:1000, Sigma), RRBP1/p180 (rabbit polyclonal, 1:1000, Sigma), PABP (rabbit polyclonal, 1:1000, Abcam), PFKB2 (mouse monoclonal, 1:1000, Cell Signaling Technology), PFKB3 (rabbit monoclonal, 1:1000, Cell Signaling Technology), PFKL (rabbit polyclonal, 1:1000, Cell Signaling Technology), PGAM1 (mouse monoclonal, 1:1000, Cell Signaling Technology), PKM (rabbit polyclonal, 1:1000, Cell Signaling Technology), RPL4 (mouse monoclonal, 1:1000, Novus), RPS6 (rabbit polyclonal, 1:1000, Cell Signaling Technology), Phospho-S6 Kinase (rabbit polyclonal, 1:1000, Cell Signaling Technology), Sec61α (rabbit polyclonal, 1:1000, Sigma), Sec61β (rabbit polyclonal^42^, 1:5000), Trap-α (rabbit polyclonal^42^, 1:5000), and UPF1 (goat polyclonal, 1:1000, Abcam).

### Mass Spectrometry

Protein samples were separated by electrophoresis on a 10 % SDS-PAGE gels and stained with Coomassie brilliant blue staining (BioBasic). Each lane on the gel was cut into 16 samples and destained using 50% acetonitrile with 0.2% ammonium bicarbonate. The gel pieces were shrunk with acetonitrile and air dried. The gel pieces were incubated in digestion mixture of trypsin (10ng/ul) in 50mM ammonium bicarbonate and incubated at 37 C overnight. The following day the gel pieces were centrifuged and the digestion mixture removed. The gel pieces were soaked in 1% formic acid in 50% acetonitrile: 49% water for 15mins, centrifuged and the supernatant combined with the digestion mixture. The samples were lyophilized and resuspended in 50mM ammonium bicarbonate and analyzed by liquid chromatograpy-tandem MS using either an LTQ-XL linear ion-trap mass spectrometer (Thermo Fisher Scientific) or Proxeon Easy-nLC 1200 pump in-line with a hybrid LTQ-Orbitrap velos mass spectromter (Thermo Fisher Scientific). Raw files from LTQ-XL mass spectrometer were uploaded to Prohits database^43^ and converted to MGF. The data was analyzed and searched using Mascot (2.3.02; Matrix Science). Raw files from LTQ-Orbitrap mass spectrometer was uploaded to Prohits and convereted to mzXML. The data was analyzed and searched using X!Tandem^44^.

We analyzed ER and cytosolic fractions from five crude polysomes preparations (both RNA-bound and ribosome-bound fractions) and two oligo-dT affinity purifications by mass spectrometry analysis. Note that due to the lower overall levels of protein in the Mock Bead pulldowns (Figure 1D), the peptide counts in this fraction are likely to be over-sampled in comparison to the oligo-dT purified samples. We scored proteins as present in the ribosome/RNA-bound fractions if represented by peptides in at least two separate experiments. For the oligo-dT experiments, proteins had to contain peptides that appeared in both experiments and be at least two-fold enriched over the mock bead pulldown in both experiments. Manual curation removed 30 proteins that were likely contaminants (e.g., Keratin, RNase, Albumin, mitochondrial proteins). Percent ER was calculated by determining the fractional representation of peptides from a protein in a given pool (pools included: ER RNA-bound, Cyto RNA-bound, ER Ribosome-bound, Cyto Ribosome-bound, ER mRNP(oligo-dT)-bound, Cyto mRNP(oligo-dT)-bound) and averaging these between experiments.

### eCLIP sequencing library preparation

Enhanced crosslinking and immunoprecipitation (eCLIP) was performed on HepG2 cells in biological duplicates, essentially as described^45^. For each replicate, extract from 20 x 106 UV-crosslinked (254 nm, 400 mJ/cm2) HepG2 cells was prepared by sonication in 1 ml lysis buffer, treated with RNase I (40 U, LifeTech), and immunoprecipitated overnight at 4 °C with 2 µg affinity-purified rabbit anti-PKM2 antibody raised against a C-terminal peptide (Sigma cat. # SAB4200105, Lot #030M4874) pre-coupled to Dynabeads sheep M-280 anti-rabbit IgG beads (LifeTech). Prior to IP, a 20 µl aliquot of extract was removed and stored at 4 °C for preparation of the size-matched input (SMInput) control. After washing, immunoprecipitated protein-RNA complexes were dephosphorylated and 3’-linker ligatated on-bead to a custom oligonucleotide adapter. All samples (IPs and SMInputs) were heated in LDS Sample Buffer (LifeTech, 70°, 10 min) and run on 4-12% NuPAGE polyacrylamide gels in MOPS running buffer (LifeTech). Complexes were wet-transferred to iBLOT nitrocellulose membranes in NuPAGE transfer buffer (all LifeTech) containing 10% methanol (overnight, 4 °C, 30V). Immunoprecipitation was confirmed by performing standard immunoblotting on a fraction of all samples. RNA-protein complexes in the range from 75 kDa (the apparent molecular mass of PKM2) to 135 kDa (corresponding to PKM2-crosslinked RNAs of ∼220 nucleotides in length) were excised from the membrane. RNA was released by proteinase K treatment in urea and recovered by phenol chloroform extraction and column purification (RNA Clean-Up kit; Zymo Research). Input samples were dephosphorylated and 3’-linker ligated to a custom oligonucleotide primer and all samples (IPs and SMInputs) reverse transcribed using AffinityScript reverse transcriptase (Agilent). After treatment with ExoSAP-IT (Affymetrix) and alkali, cDNAs were recovered by purification on Dynabeads MyONE Silane beads (LifeTech), 5’-linker ligated on-bead to a custom oligonucleotide primer, purified, and recovered in 27 µl. cDNA was quantified by qPCR analysis of a fraction each sample using oligonucleotide primers specific to the 5’ and 3’ adapters. Half of the recovered cDNA was PCR-amplified (Q5 polymerase, NEB) using custom sequence-indexed oligonucleotide primers with the following cycle numbers: replicate 1: input 8, IP 15; replicate 2: input 8, IP 13. PCR products were purified (Agencourt AMPure XP beads; Beckman Coulter), size-selected to 175 −350 bp on 2% agarose gels, extracted (MinElute Gel Extraction kit; Qiagen), and quantified on a TapeStation using D1000 ScreenTape (Agilent). Libraries were sequenced on a HiSeq 4000 instrument (Illumina) in paired-end 55 bp mode.

### Computational analysis of eCLIP-Seq data

#### Trimming and mapping

Sequencing reads were processed essentially as described^45^. Reads were adapter-trimmed and mapped to human-specific repetitive elements from RepBase (version 18.04) by STAR^46^. Repeat-mapping reads were removed and remaining reads mapped to the human genome assembly hg19 with STAR. PCR duplicate reads were removed using the unique molecular identifier (UMI) sequences in the 5’ adapter and remaining reads retained as ‘usable reads’. Peaks were called on the usable reads by CLIPper^47^ and assigned to gene regions annotated in Gencode (v19). Each peak was normalized to the size-matched input (SMInput) by calculating the fraction of the number of usable reads from immunoprecipitation to that of the usable reads from the SMInput. Peaks were deemed significant at > 8-fold enrichment and p-value < 10^-5^ (Chi-square test). All sequencing and processing statistics are in Supplemental Table 3.

#### De novo motif analysis

HOMER^48^ was used to identify de novo motifs using the command ‘findMotifs.pl <foreground> <sequence database> <output location> -rna -bg <background>’. The sequence database that was used corresponded to coding sequences of genes downloaded from Ensembl v96^49^: for each gene, we selected the coding sequence of the principal isoform, as defined by APPRIS^50^, followed by masking the endogenous repeat elements using RepeatMasker (http://www.repeatmasker.org) to ensure that motif finding was not confounded by sequence similarity of these elements. The foreground consisted of the genes that we found to be PKM- and glucose-sensitive and were also bound by PKM, whereas the background consisted of all the expressed genes. We also repeated the same analysis separately for genes that were PKM-regulated, glucose-regulated, or were bound by PKM (as defined by eCLIP), and used MoSBAT^51^ to identify the highest-scoring motif that was commonly found in all these analyses.

### Radiolabeling of PKM2-bound RNA fragments

Immunoprecipitation was End-labeling was performed as described^52^ with modifications. 20 million UV-crosslinked HepG2 cells were lysed in 550 µl lysis buffer (50 mM Tris-HCl pH 7.4, 100 mM NaCl, 1% NP-40, 0.1% SDS, 0.5% sodium deoxycholate) with protease inhibitor cocktail (Roche). Lysates were sonicated for 5 minutes (BioRuptor, low setting, 30 s on/off) in an ice-cold water bath. After addition of 2.2 µl Turbo DNase (NEB) and 5.5 µl RNase A (Millipore cat. #20-297), diluted 1:100 (high RNase) or 1:10,000 (low RNase) in low-stringency wash buffer (20 mM Tris-HCl, pH 7.4, 10 mM MgCl2, 0.2% Tween-20), samples were incubated at 37 °C for 5 minutes with shaking. RNase digestion was stopped with 11 µl Murine RNase inhibitor (NEB) and insoluble material removed by centrifugation (15 min, 15,000 g, 4 °C). Protein-RNA complexes were immunoprecipitated overnight at 4 °C with PKM2 antibody (SAB4200105, Sigma) or normal rabbit IgG (Thermo Fisher) pre-coupled to magnetic beads (Dynabeads M-280 Sheep anti-Rabbit IgG, Thermo Fisher). A series of wash steps was employed to ensure stringency, as follows: 2x low stringency wash buffer (see above), 2x high stringency buffer (15 mM Tris-HCl pH 7.4, 5mM EDTA, 2.5 mM EGTA, 1% Triton-X 100, 1% sodium deoxycholate, 0.1% SDS, 120 mM NaCl, 25 mM KCl), 2x high salt wash buffer (50 mM Tris-HCl, pH 7.4, 1M NaCl, 1mM EDTA, 1% NP-40, 0.1% SDS, 0.5% sodium deoxycholate), 2x low stringency wash buffer, and 2x PNK buffer (50 mM Tris-HCl pH 7.4, 10mM MgCl2, 0.5% NP-40). Protein-RNA complexes were radiolabeled on-bead in 40 µl reactions with T4 polynucleotide kinase (NEB) and 2 µl [γ-32P]ATP (6,000 Ci/mmol, 10 mCi/ml) for 10 minutes at 37 °C. Beads were washed 3x in PNK buffer, resuspended in NuPAGE LDS Sample Buffer (Thermo Fisher) containing 0.1 M DTT. Protein-RNA complexes were denatured at 75 °C for 15 mins and run on 4%–12% NuPAGE Bis-Tris gels in NuPAGE MOPS running buffer (all Thermo Fisher) at 150V for 1.5 h, wet-transferred to nitrocellulose membrane using NuPAGE transfer buffer (Thermo Fisher) with 10% methanol for 3h at 200 mA. The membrane was exposed to film for 20 minutes at room temperature and the film developed.

## Acknowledgements

We would like to thank A. Cochrane, N. Sonenberg and J. Cate for generously sharing various antibodies, J. Rini and D. Zhou for providing 293F-HEK cells and J. Glover, C. Smibert and SickKids’ core facility for the use of equipment. We also thank E. Lee, P. Ho, H. Zhang, for experimental help. We also thank C. Smibert, H. Wyatt, T. Moraes and A. McQuibban for comments on the manuscript. A.F.P. was supported by funding from the Canadian Institutes of Health Research and The Natural Sciences and Engineering Research Council of Canada.

---

**Supplemental Figure 1.**
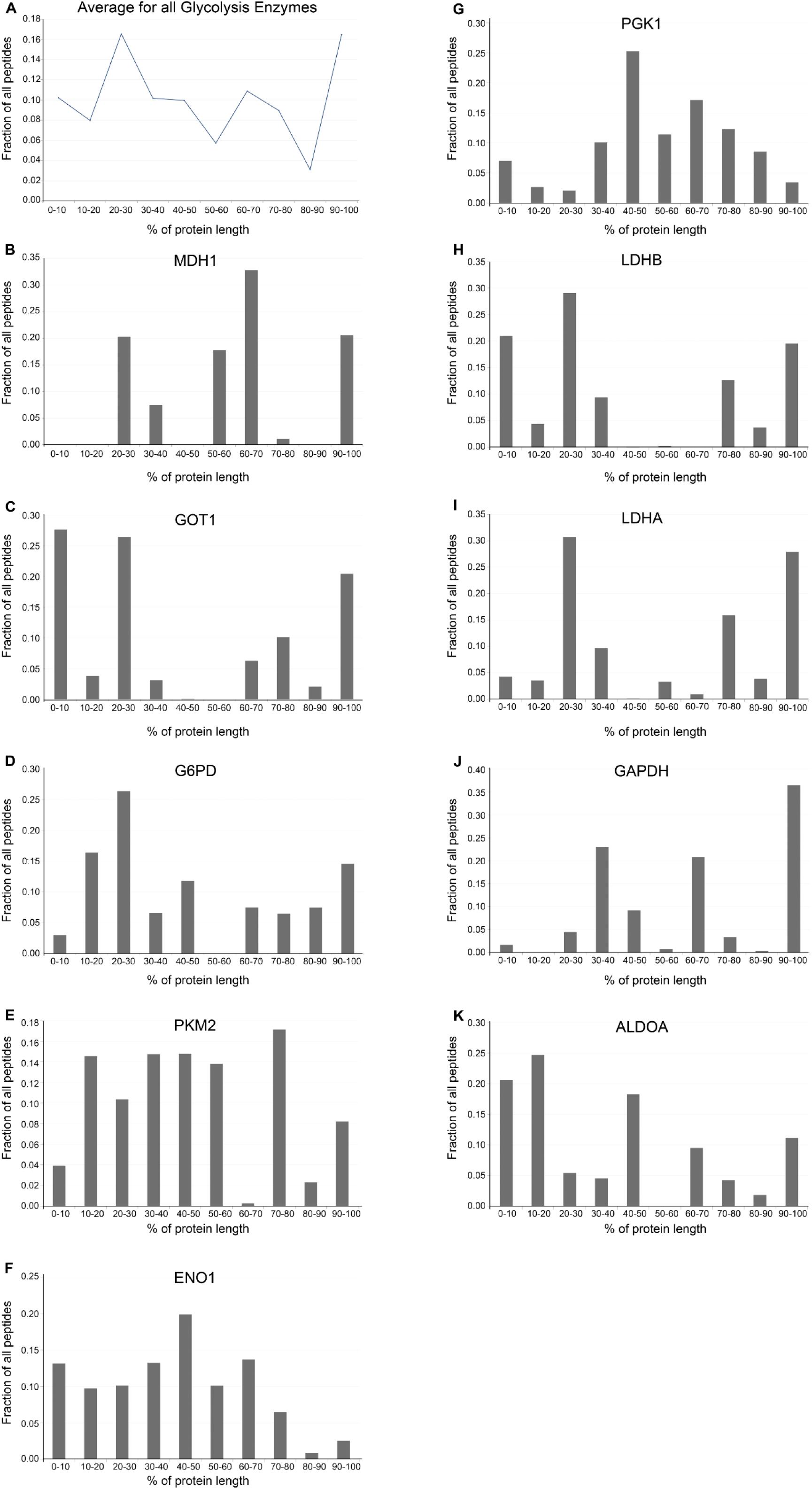
Distribution of peptides from glycolytic proteins isolated by oligo-dT affinity chromatography. For all the listed proteins, the number of spectral counts for peptides along the length of the protein were compiled and binned. The distribution of the fraction of total peptides (*y-axis*) along the protein length for each bin (from N-to C-terminus; *x-axis*) were plotted. A) The overall average for all glycolysis enzymes. Note the large fraction of peptides that map back to the C-termini (bin 90-100%). B-K) The distribution for individual genes. Note that with the exception of ENO1 and PGK1, a significant of peptides isolated in the oligo-dT affinity purifications map back to the C-termini (bin 90-100%).

**Supplemental Figure 2.**
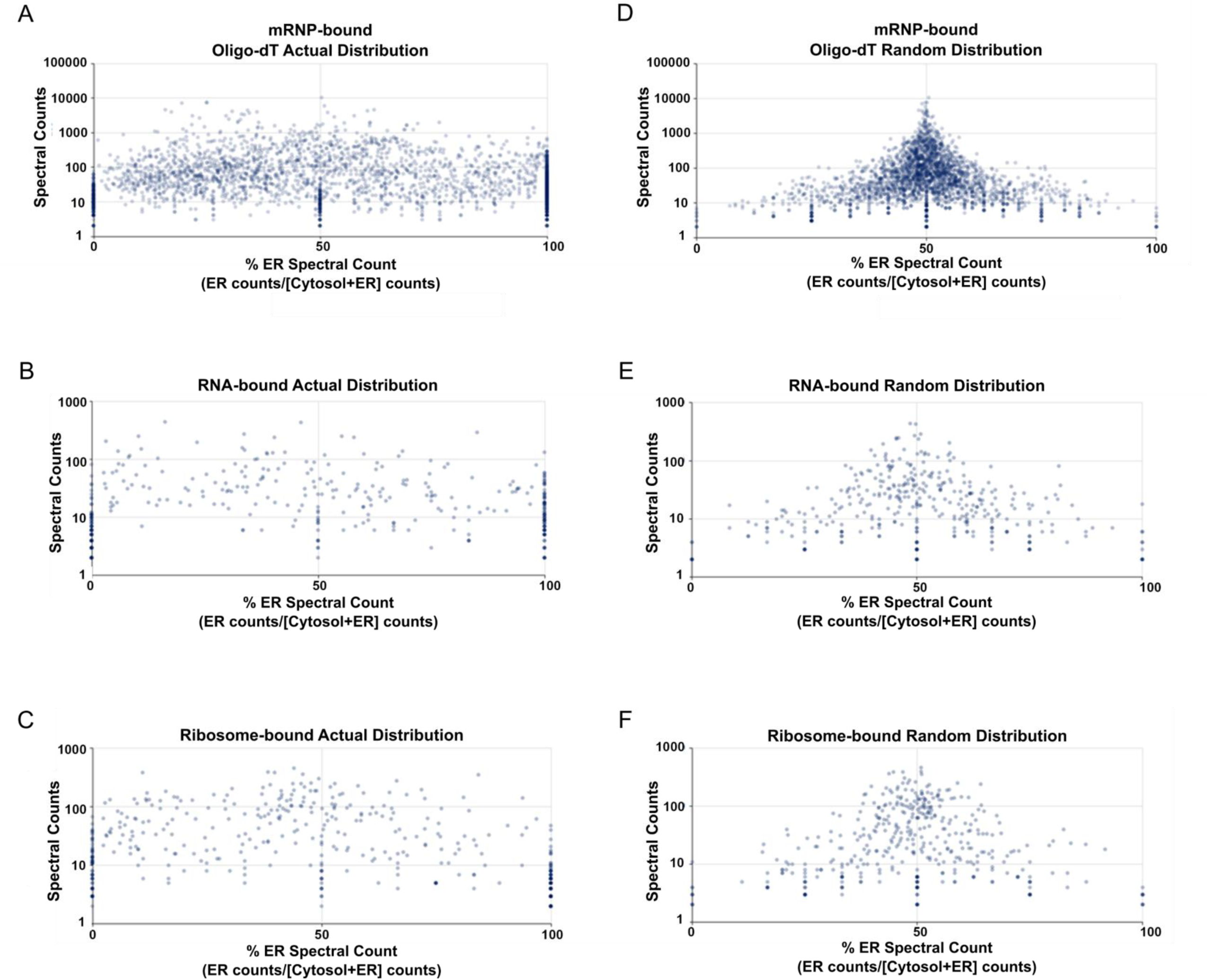
The distribution of many polysome-associated proteins differs significantly between the ER and cytosol. A-C) For each protein present in both crude polysome and oligo-dT purifications in our list of mRNA/Ribosome-associated proteins (see Supplemental Table 1), the total number of peptides (*y-axis*) identified by mass spectrometry was plotted against the percentage of peptides found in the ER [100% x peptides in ER/(peptides in ER + peptides in cytoplasm); ‘% on ER’ - *x-axis*]. Note that data in (A-C) is identical to that of Figure 1J-L. D-F) The same as (A-C) except that each peptide was randomly reassigned to either the ER or cytosolic fraction and the ‘% on ER’ was recalculated.

**Supplemental Figure 3.**
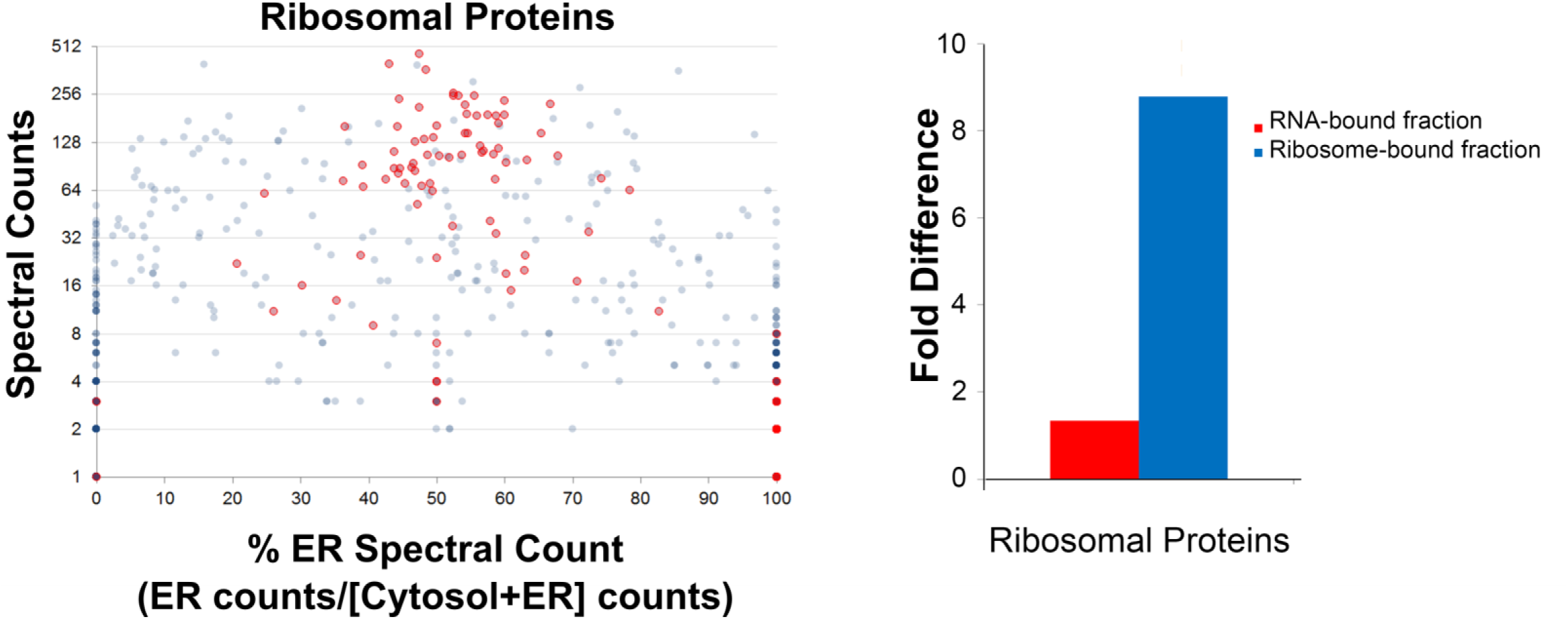
Ribosomal proteins were equally sampled between the ER and cytosol. A) For each protein in our list of polysome-associated proteins (see Supplemental Table 1), the total number of peptides (*y-axis*) identified by mass spectrometry was plotted against the percentage of peptides found in the ER [100% x peptides in ER/(peptides in ER + peptides in cytoplasm); ‘% on ER’ - *x-axis*] for the ribosome-bound fractions. This data is identical to that of Figure 1K. Note that the ribosomal proteins, which are labeled in red, are the most abundant proteins and are equally sampled between the ER and cytosol. B) The normalized number of ribosomal peptides in the RNA-bound fraction versus the ribosome-bound fraction. Overall ribosomal proteins made up only 5% of the peptides in the RNA-bound fractions and 22% of the peptides from the ribosome-bound fractions.

**Supplemental Figure 4.**
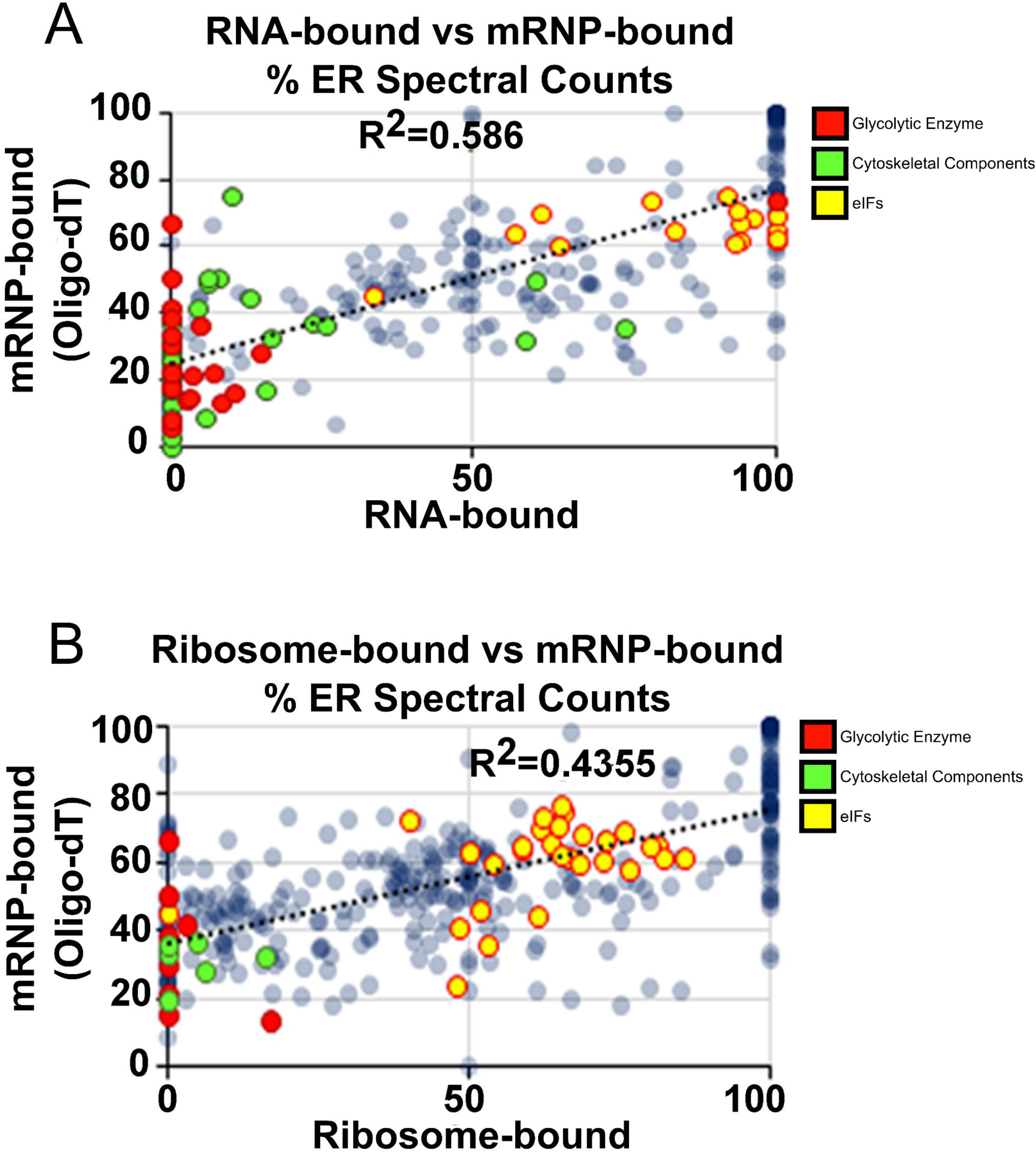
ER/Cytosolic Distribution of RNA and Polysome-Associated Proteins. Correlation between the percentage of peptides found in the ER/total in the RNA-bound fraction of the crude polysome preparation and in the oligo dT-associated proteins (A) and in the ribosomal-bound fraction of the crude polysome preparation and in the oligo dT-associated proteins (B). The data is identical to that of Figure 1J-L but replotted to compare the various fractions. Classes of proteins that are enriched in either the ER or the cytoplasm are highlighted as in Figure 1J-L – carbohydrate metabolic proteins (red), eIFs (yellow), and cytoskeletal-associated proteins (green).

**Supplemental Figure 5.**
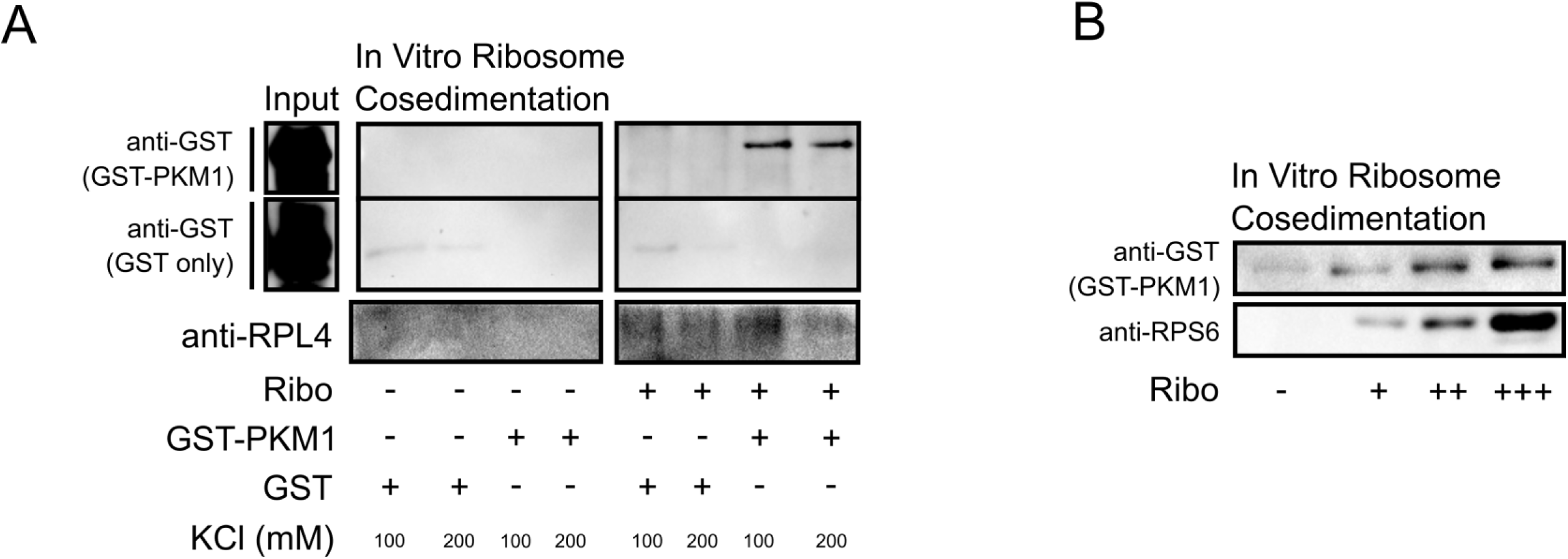
Association of PKM with salt-washed ribosomes. A) Bacterial-expressed GST or PKM1-GST were incubated with and without salt-washed ribosomes, isolated from HEK293T cells. Incubated reactions were sedimented through a sucrose cushion and were immunoprobed for GST or RPL4. B) Bacterial-expressed PKM1-GST was incubated with increasing amounts of salt-washed ribosomes and then sedimented through a sucrose cushion. Note that the amount of PKM1-GST in the pellet rises with increasing levels of ribosomes.

**Supplemental Figure 6.**
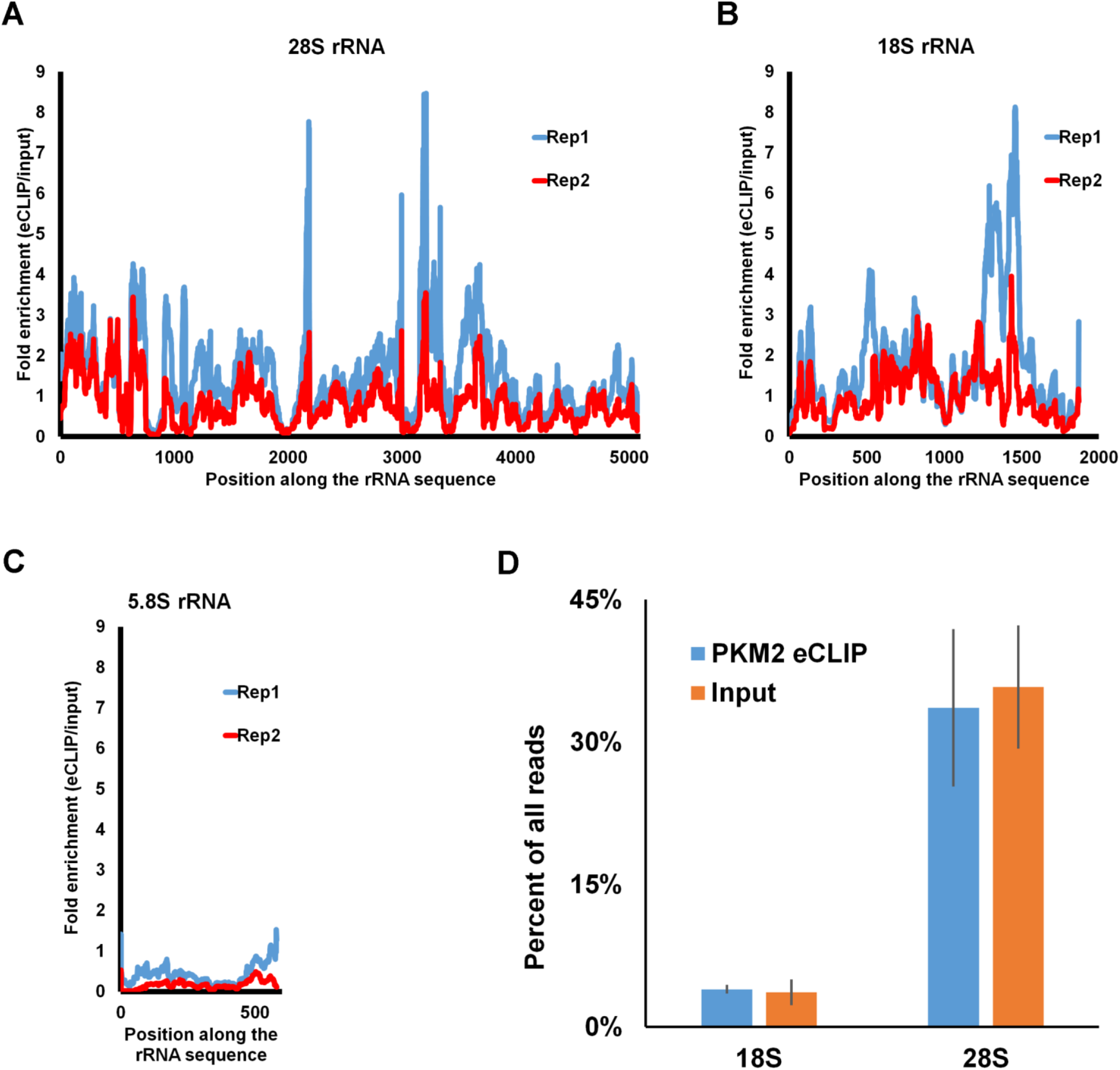
PKM2 eCLIP peaks mapping to rRNA. A-C) PKM2 eCLIP reads were compared to size-matched input reads using RNA repeat-family centric mapping (Van Nostrand et al., manuscript in prep.). The densities of eCLIP/input reads (*y-axis*) was plotted along the length (*x-axis*) of the 28S, 18S and 5.8S rRNA genes. D) The fraction of PKM2 eCLIP and input reads vs total reads were plotted. Each bar represents the average and standard deviation of the two replicates.

**Supplemental Figure 7.**
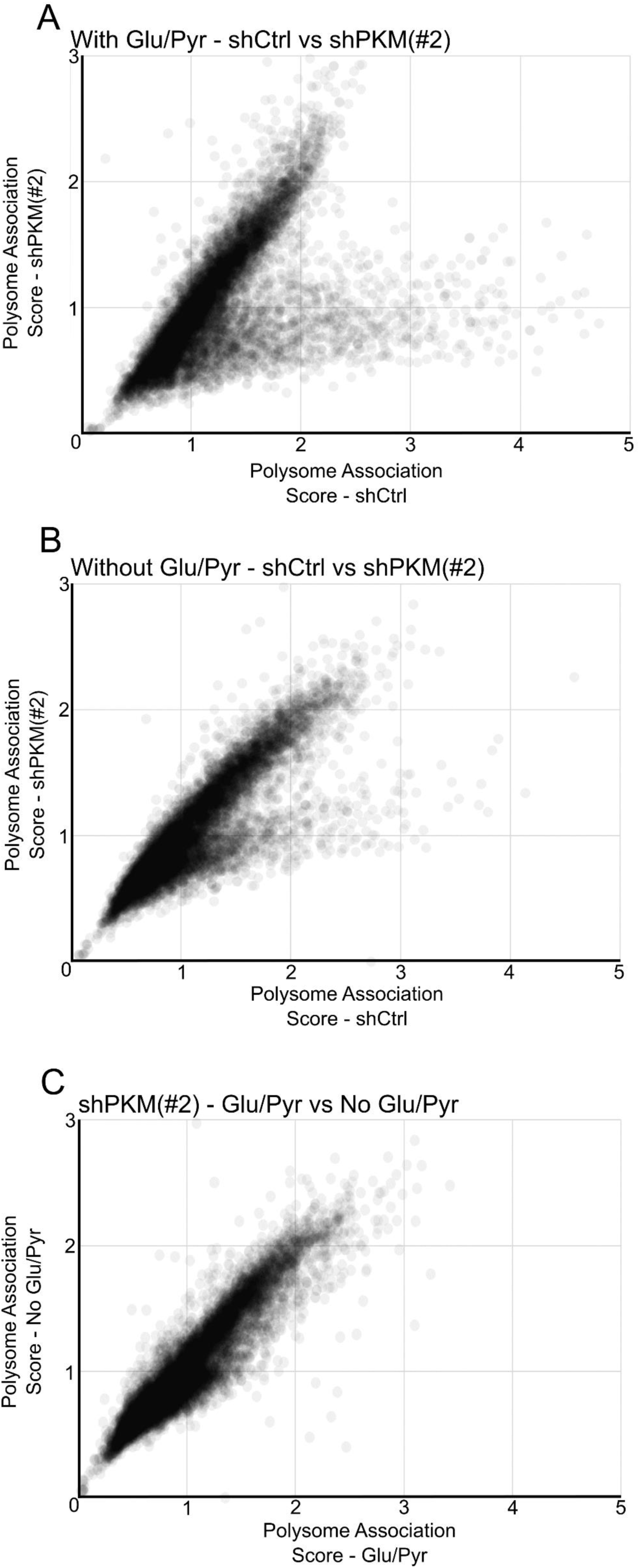
PKM inhibits the translation elongation of ribosomes on a subset of mRNAs in a glucose/pyruvate-dependent manner. Analysis performed as in Figure 4C and D except using a different lentiviral generated shRNA against PKM.

**Supplemental Figure 8.**
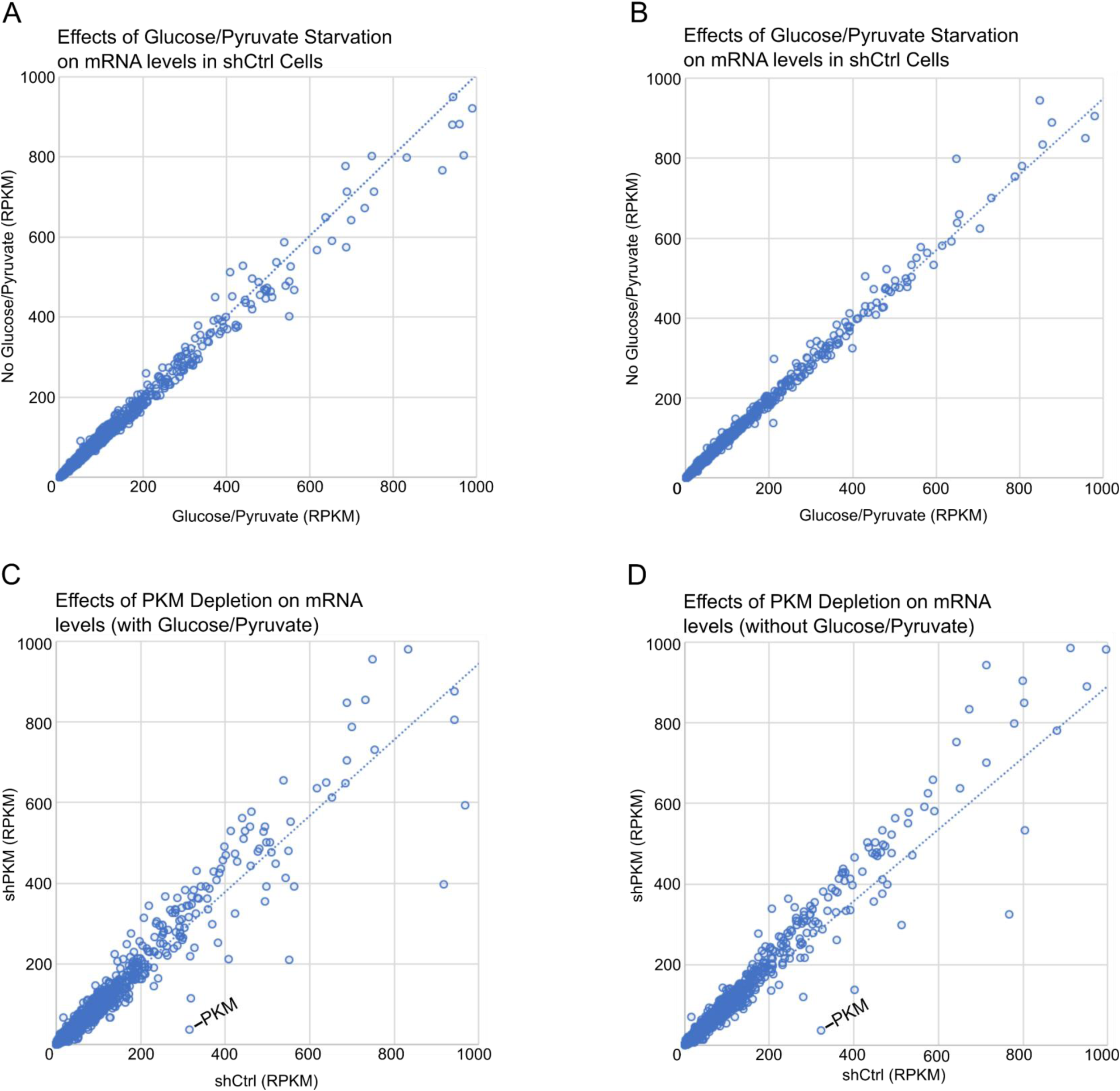
PKM-depletion and glucose/pyruvate starvation do not drastically effect steady state mRNA levels. A-D) A comparison of RPKM of detected genes from the input of control or PKM-depleted U2OS cells incubated in either DMEM with or without glucose/pyruvate for 3 hours.

**Supplemental Table 1. Mass Spectrometry of mRNA/Ribosome-bound proteins**

The list of 496 mRNA/Ribosome-bound proteins. Proteins must have peptides that appear in 1) at least two separate RNA-bound or Ribosome-bound experiments (out of a total of five); and 2) in both mRNA-bound (oligo-dT affinity chromatography) experiments and have at least two fold more peptides in the oligo-dT affinity chromatography precipitates than in the mock bead precipitates in both experiments. Each spectral count column represents the total counts for all experiments. Each percent ER column represents the average of at least 2 experiments. The percent ER is calculated by the fractional representation of spectral peptide counts for a given protein in the ER for each fraction (RNA-bound; Ribosome-bound; mRNP-bound, see methods) divided by the fractional representation of spectral counts for the same protein in the ER and cytosol combined for that same experiment.

**Supplemental Table 2 RBPome Comparisons**

**Supplemental Table 3. PKM eCLIP Statistics**

**Supplemental Table 4. PKM2 eCLIP Peaks**

**Supplemental Table 5. Glucose and PKM-Regulated translation**

The translation of an mRNAs was scored as being sensitive to PKM-depletion if 1) the change in polysome-association (reads in polysome/reads in input) dropped by at least two fold between the two shRNA experiments for the glucose-treated conditions (control-depleted vs PKM-depleted) and 2) that the drop in polysome association was greater than three standard deviations for the glucose-treated conditions (control-depleted vs PKM-depleted). The translation of an mRNAs was scored as being sensitive to glucose if the change in polysome-association (reads in polysome/reads in input) dropped by at least 50% between the glucose/pyruvate-treated and -starved conditions for the control-depleted cells.

